# Dynamics of the rice yellow mottle disease in western Burkina Faso: epidemic monitoring, spatio-temporal variation of viral diversity and pathogenicity in a disease hotspot

**DOI:** 10.1101/2023.03.27.534376

**Authors:** Estelle Billard, Mariam Barro, Drissa Sérémé, Martine Bangratz, Issa Wonni, Moustapha Koala, Abalo Itolou Kassankogno, Eugénie Hébrard, Gaël Thébaud, Christophe Brugidou, Nils Poulicard, Charlotte Tollenaere

## Abstract

The rice yellow mottle virus (RYMV) is a model in plant virus molecular epidemiology and phylogeography, with the reconstruction of historical introduction routes at the scale of the African continent. However, information on patterns of viral prevalence and viral diversity over multiple years at local scale remain scarce, in spite of potential implications for crop protection.

Here we describe a five-years monitoring of RYMV prevalence in six sites from western Burkina Faso. This study confirmed one irrigated site as a disease hotspot, and found two rainfed lowland sites with occasional high prevalence levels. Within studied field, a pattern of disease aggregation was evidenced at a five-meter distance, as expected for a mechanically transmitted virus.

Next, we monitored RYMV genetic diversity in the irrigated disease hotspot site, revealing a high viral diversity, with the current coexistence of various distinct genetic groups at the site-scale (irrigated perimeter of ca. 520 ha), and also within various specific fields (25 meters side). One genetic lineage, named S1bzn, is the most recently introduced group and increased in frequency over the studied period. Its genome results from a recombination between two other lineages.

Finally, experimental work evidenced no differences between three rice varieties cultivated in Burkina Faso in terms of resistance level, and no statistical effect of RYMV genetic group on symptom expression and viral load. We found however, that infection outcome depended on the specific RYMV isolate, with various isolates from the lineage S1bzn found to be particularly aggressive, including one accumulating at highest level.

Overall, this study documents a case of high viral prevalence and high viral diversity, with the co-occurrence of divergent genetic lineages at small geographic scale. A recently introduced lineage, that includes viral isolates with high symptoms and accumulation in controlled conditions, could be recently rising though natural selection. Following up the monitoring of RYMV genetic and pathogenic diversity in the area is required to confirm this trend and further understand the factors driving the maintenance of viral diversity at local scale.

## INTRODUCTION

Biotic constraints drastically affect crop production, with pathogens and pests being responsible for an estimated 17-30% of global yield losses (Savary et al., 2019). Food-deficit regions with fast-growing populations are disproportionately affected, notably Sub-Saharan Africa (Savary et al., 2019), where production losses were estimated to reach over one billion dollars and threaten food security (Sileshi & Gebeyehu, 2021). Viruses cause almost half of the reported plant emerging infectious diseases (Anderson et al., 2004), and constitute a major challenge to agriculture, with an annual global cost of more than 30 billion dollars (Sastry and Zitter, 2014). Such virus-induced yield losses may even increase in the next decades, as a consequence of climate change and agricultural intensification, urging to improve the scientific understanding of epidemiological dynamics of crop viral diseases (Jones & Naidu, 2019).

Spatio-temporal studies of crop virus dynamics are of major importance to better understand the drivers of epidemics and to develop appropriate control strategies (McLeish, Fraile & García-Arenal, 2021). In this respect, several studies have successfully used knowledge of small-scale crop virus dynamics to guide disease management methods, as exemplified by the long-term study of viruses infecting cucurbit and solanaceous crops and weeds in southern France (Desbiez et al., 2009, 2020). Virus detection, genetic and pathogenic diversity (including in neighboring weeds) and spatial structure analyses, inform for each considered virus the respective key prophylactic and cultural measures to undertake: control of insect vectors, weeding, deployment of specific resistance genes and sanitation of human-assisted plant material exchanges (Desbiez et al., 2009, 2020).

For two tomato begomoviruses in Brazil as well, Macedo et al. (2019), revealed that primary spread was the most important mechanism for epidemics of the two common whitefly-transmitted viruses, so that preventing the arrival of viruliferous insects to production areas is the key to manage golden mosaic and yellowing diseases. Finally, a 21-years monitoring of pepper-infecting tobamoviruses in southeastern Spain allowed to document the rapid genetic viral diversification related to the deployment of pepper resistant cultivars (Fraile et al., 2011). Overall, spatial and temporal variation in prevalence and diversity allow to identify disease hotspot, inoculum sources and origin of new pathogen variants, to characterize transmission roads, and identify newly adapted pathogen genotypes that are likely to increase in frequency and spread. All this information is mandatory for science-based deployment of control methods to efficiently reduce virus epidemics and emergence risks (McDonald & Linde, 2002; Burdon & Thrall, 2008; Picard et al., 2017; Desbiez et al., 2020; Saubin et al., 2022).

Rice (*Oryza spp.*) is of critical import regarding present and future food security, with more than 3.5 billion people now relying on it for more than 20% of their daily diet (Ahmadi & Bouman, 2013). In West Africa, rice consumption experienced a recent 8% increase each year between 2009 and 2019 (Soullier et al., 2020). To face this growing demand, proactive policies led to a rapid increase in rice production with an average annual growth higher than 10% between 2009 and 2019 (Soullier et al., 2020). This increase in rice production is the consequence of increased areas cultivated with rice, and agricultural intensification. However, intensification of rice cropping may render rice production more vulnerable to disease and pest outbreaks in West Africa. While as much as 16 viruses were reported worldwide (Compendium of rice diseases and pests; Cartwright et al., 2018), only two rice viruses have been reported to date in West Africa: *Rice stripe necrosis virus* (RSNV) and, most importantly in terms of yield impact, *Rice yellow mottle virus* (RYMV). RYMV is a *Sobemovirus* (family *Solemoviridae*) endemic of Africa (see Hébrard et al., 2021 for review), mechanically transmitted either by insects or other animals, or during human-mediated agricultural manipulations (Traoré et al., 2009). RYMV causes severe yield losses to rice production (Hébrard et al., 2021), and was consequently listed within the 14 emerging infectious pathogens threatening food security (Savary et al., 2019) and economies in Africa (Sileshi & Gebeyehu, 2021), and in the top ten list of economically important plant viruses (Rybicki, 2015).

RYMV is also one of the major model for plant virus molecular epidemiology (Trovão et al., 2015; Pagán, Fraile & García-Arenal, 2016). Continental-scale molecular epidemiology studies were undertaken (Pinel-Galzi et al., 2009, 2015), and compared to RYMV field epidemiology (Rakotomalala et al., 2019; Issaka et al., 2021), revealing a strong geographic structuration of genetic diversity (Pinel-Galzi et al., 2015). Various viral strains may however overlap in some regions, as exemplified by the coexistence of several RYMV strains in Côte d’Ivoire (N’Guessan et al., 2000), or at the local scale in Tanzania (Kanyeka et al., 2007). Description of recombinant isolates in East Africa (Pinel-Galzi et al., 2009; Ochola et al., 2015; Adego et al., 2018) suggests a substantial level of viral strain co-occurrence and co-infection, but this remains poorly documented. Indeed, compared to country or continental-scale studies, few studies described local-scale RYMV epidemics in terms of disease prevalence and genetic diversity. Such local monitoring is however the relevant scale for the efficient deployment of resistant cultivars.

In Burkina Faso, yellow mottle disease was first reported in 1981 (John, Thottapilly & Awoderu, 1984; Salaudeen et al., 2010) and research on this virus began with an intensive sampling in 1994-1995 (Konate, Traore & Coulibaly, 1997). Yellow mottle disease was known by the farmers from western Burkina Faso: reported as the most serious rice disease by farmers from ten villages of the Cascades Region (in southwestern Burkina Faso, Kam et al., 2013) and as a major concern for rice cultivation in the irrigated perimeter of Banzon (Traoré et al., 2015). Burkina Faso is located at the crossroads of RYMV dispersal roads in West Africa (Dellicour et al., 2018), and is one of the countries where various RYMV strains have been reported (S1wa, S1ca, S2; Pinel-Galzi et al., 2015), with potential contrasting resistance-breaking capacities (considering the S1ca pathotype T’ able to evolve and infect resistant cultivars under controlled conditions and reported in border countries; Hébrard et al., 2018). In western Burkina Faso, Barro et al. (2021a) monitored four major rice diseases in six study sites (three irrigated areas and neighboring rainfed lowlands) over four years (2016-2019), following an initial survey of viral and bacterial diseases in a selection of sites in 2015 (Tollenaere et al., 2017). Over the 2016-2019 period, yellow mottle disease symptoms were observed in 54 fields over the 179 observations (30.2% of visited rice fields). The interaction between rice production system (irrigated area vs rainfed lowland) and the geographical zone (three zones distant from each other from 40 to 90 km) was highly significant to explain yellow mottle occurrence. Indeed, the irrigated area of Banzon and, to a lesser extent, the rainfed lowland of Bama (Badala village) were identified as yellow mottle disease hotspots, while the four other sites presented much fewer symptomatic plants (Barro et al., 2021a). Banzon is a 520-ha irrigated perimeter located approximately 60 km west of the city of Bobo-Dioulasso. As a consequence of the creation of the irrigation scheme (initiated in 1973 and finished in 1981), the population has increased tenfold from 1975 to 1985, and the practice of fallow disappeared (Toe, 1992).

The present study complements the previous study of multiple rice diseases in western Burkina Faso by Barro et al. (2021a), and focuses specifically on the rice yellow mottle disease, with the following questions that have never been addressed for RYMV: What is the spatio-temporal dynamics of the disease at a regional scale and can we identify drivers of infection at local (field) and regional scales? What is the level of viral genetic diversity at the local scale and does it vary through time? Do various viral strains co-exist within the same irrigated area or even the same field? If so, do these strains differ in terms of pathogenicity on susceptible rice host? Globally, this study aims to monitor the dynamics of epidemics and characterize the drivers of infection and genetic diversity for the major viral threat of rice, a staple food far from self-sufficiency in West Africa.

## MATERIAL AND METHODS

### Longitudinal sampling in selected farmer’s fields in western Burkina Faso

Symptom observations and sampling were performed between 2015 and 2019 in six sites of western Burkina Faso (Bobo-Dioulasso area). Only two of these sites (Karfiguela and Banzon) were studied in 2015. Then, annually from 2016 to 2019, we visited three irrigated areas and three rainfed lowlands located nearby the irrigated sampled sites, resulting in six sites in total, see Barro et al. (2021a). These visits were always performed at the maximum tillering, or heading initiation, growth stage (September-early December). In each selected field (25x25m), a 4x4 grid was delimited and symptoms associated with multiple pathogens were assessed visually in the four cells of the diagonal of the grid (Barro et al., 2021a). Yellow mottle symptoms were scored from 0 (no symptom observed) to 100% (all plants symptomatic) in the four diagonal cells of each studied field. In addition, the 16 plants located at the grid nodes were inspected for symptoms and sampled (three leaves put in an envelope and dried using silica gel). In every case, we obtained permission from the farmers to work (observations and leaves sampling) in their fields. We interviewed most farmers to get data on the cultivar and agricultural practices for each studied field, as we also asked questions to document constraints to rice production, farmer’s knowledge of rice diseases and control methods. These farmer’s interview data were analyzed in Barro et al. (2021a) mostly to contrast the major agricultural practices used in each rice growing system. Further details on the methodology, and all the data (symptom observation as well as farmer’s interviews) for each field studied every year, are available in DataSuds: https://doi.org/10.23708/8FDWIE.

### Enzyme-linked immunosorbent assay (ELISA) detection of RYMV in collected samples

In all cases of observation of specific yellow mottle disease symptoms in a field, the 16 sampled plants were analyzed by RYMV-specific serological detection. Each plant sample consisted on one 1-cm long leaf sample from each of the three different leaves sampled (3-cm equivalent). Serological detection was performed on these samples, using a previously described DAS-ELISA (Double Antibody Sandwich Enzyme Linked Immunosorbent Assay) protocol with the RYMV-Mg antiserum (Ndjiondjop et al., 1999; Traoré et al., 2008). Absorbance at 405 nm was read using a Metertech Σ960 microplate reader in LMI PathoBios detection platform (Kamboinse, Burkina Faso) for 2015-2017 samples or a Tecan Spark Multimode plate Reader in IRD-Genetrop platform (Montpellier, France) for 2018-2019 samples. Following Traoré et al. (2008), samples were considered as positive if the optical density (OD) obtained was higher than the average of negative samples plus three times the standard deviation. A constant threshold of OD =0.3 was also used to compare the obtained results to a more conservative analysis (see also N’Guessan et al., 2000).

### RYMV-specific ELISA data analysis

We used the R software (R Core Team, 2018) and the package *ggplot2* (Wickham, 2016) to visualize prevalence data (frequency of infection within fields) in the different sites and years. Generalized Linear Models (GLMs) were used to test the effect of the variable ‘year’ and the site (included in the model as the interaction between ‘rice system’ and ‘zone’) on RYMV prevalence based on serological detection (binomial variable), either by including or not the fields were no yellow mottle symptoms were observed (prevalence =0). Post hoc tests were performed using the package *lsmeans* (Lenth, 2016). In addition, the relationship between prevalence estimates based on DAS-ELISA data and symptom-based prevalence data was tested using a Spearman correlation test (*cor.test* function).

We then tested for an effect of agricultural practices on yellow mottle disease prevalence. Because previous analyses revealed a very strong effect of the site, we performed this analysis on a subset of the data: all observations performed within the irrigated perimeter of Banzon (Banzon-IR), revealed as a yellow mottle disease hotspot by the above-mentioned analyses. Three agricultural practices presented variability within the Banzon-IR dataset: rice cultivar (four cultivars), off-season cultivation (rice, or nothing), and the number of urea application (0 or 1 application, compared to 2 or more applications). However, off-season cultivation and the number of urea application were highly correlated so off-season cultivation was only kept in the model. Generalized Linear Mixed Models (GLMMs), were performed using the R package *lme4* (Bates et al., 2014), on the binomial variable of ‘RYMV prevalence’ (from DAS-ELISA data). Year was included in the model as a random effect and the agricultural variables (‘rice cultivar’ and ‘off-season cultivation’) and their interaction, as fixed explanatory variables.

Finally, we tested whether presence/absence data (from RYMV symptom observations on the one hand and from DAS-ELISA detection result on the other hand) were spatially aggregated at the field (25x25m) level. To this aim, analyzes were performed using the same methodology as described in Thébaud et al. (2005). The test compares the observed distribution of distances between infected plants to the simulation under *H_0_* (random distribution of infected plants, i.e. no spatial aggregation). This test was performed over the fields presenting at least two infected plants (29 cases for the symptoms and 33 cases for DAS-ELISA).

### Sanger sequencing of Coat Protein (CP) gene and viral genome

Genetic analysis of collected samples was performed in the irrigated area of Banzon only, and particularly for samples collected within five fields (distant from ca. 400 m to 2.2 km), chosen for their high RYMV prevalences over the five sampling years.

Amplification of the coat protein (CP) gene (ORF4) was performed by RT-PCR as described earlier (Pinel et al., 2000; Fargette et al., 2004), using the primer RYMV-M 5’- CGCTCAACATCCTTTTCAGGGTAG-3’for reverse transcription (RT) and the primer pair RYMV- III 5’-CAAAGATGGCCAGGAA-3’ and RYMV-M for the PCR. Amplified products were sent to Genewiz (Leipzig, Germany) for Sanger sequencing with the same primer pair used for the PCR. A 720- bp sequence was obtained for 132 isolates and alignment was performed using MEGA-X software.

We performed full-length genome sequencing for a few representative isolates, using the approach described in (Fargette et al., 2004). Briefly, a RT was performed using the primer RYMV-II 5’- CTCCCCCACCCATCCCGAGAATT-3’ and two PCRs were performed with the two pairs of primers: RYMV-R1 5’-CAATTGAAGCTAGGAAAGGAG-3’ with RYMV-14bis 5’- ACTTCGCCGGTTTCGCAAAGGATT-3’; and RYMV-2136 5’-CATGCTGGGAAAAGTGTCTG-3’ with RYMV-II. Additional primers were used for sequencing using the Sanger technique (Genewiz, NJ, USA).

Obtained sequences were deposited in GenBank (Accession Nos. to be assigned).

### Genetic diversity, phylogenetic analyses and detection of recombination events

The ORF4 sequence dataset and full-length genome sequences obtained from Banzon were compared to those available in NCBI (cf. Issaka et al., 2021), *i.e.* 261 CP gene sequences and 37 complete genome sequences from isolates collected in West and Central Africa.

Multiple sequences were aligned using MUSCLE (Edgar, 2004) implemented in SEAVIEW v4.7 (Gouy, Guindon & Gascuel, 2010). We reconstructed Maximum-Likelihood (ML) phylogenetic trees with SEAVIEW using the best-fitting nucleotide substitution models (K2+G+I and GTR+G+I for the CP gene and the complete genome datasets, respectively) determined with MEGAX (Kumar et al., 2018) and 100 bootstrap replicates. Phylogenetic trees were drawn using FigTree v1.3.1 (http://tree.bio.ed.ac.uk/software/figtree/). In addition, we used the median-joining network method implemented in the Network software (version 4.611; www.fluxus-engineering.com) (Bandlet, Forster & Röhl, 1999) to reconstruct the minimum spanning network (MSN) connecting all the CP sequences (at the amino acid level) obtained in this study.

Genetic diversity of the CP gene dataset was estimated per year, field and rice cultivars using the K2+G+I substitution model, with standard errors of each measure based on 100 bootstrap replicates, as implemented in MEGAX. Difference in nucleotide diversity of the virus populations among groups (*i.e.* year, field and rice cultivars) was tested by analysis of molecular variance (AMOVA), as implemented in Arlequin v. 5.3.1.2 (Excoffier, Laval & Schneider, 2005). AMOVA calculates the *F*_ST_ index explaining the between-group fraction of total genetic diversity. Significance of these differences was obtained by performing 1000 permutations.

Potential recombination signals from complete RYMV genome sequences or CP gene dataset were searched using the pairwise homoplasy test (PHI test) using SplitsTree v4.18.3 (Huson & Bryant, 2006) and the seven algorithms implemented in the RDP program v4.97 (Martin et al., 2015). Only recombination events detected by five or more methods and with *P*-values above 10^-5^ were considered.

### Bayesian evolutionary inferences and discrete phylogeographic analyses

In order to examine the degree of temporal signal in the CP gene dataset, we first used an exploratory linear regression approach (Duchêne et al., 2015; Murray et al., 2016). Based on ML phylogenetic tree reconstructed from the CP gene dataset, the temporal signal of this sequence dataset was visualized and tested in PhyloStemS (Doizy et al., 2020) to regress phylogenetic root-to-tip distances against sampling date using the root that minimized the residual mean squares. In addition, the significance of the temporal signal was evaluated by a date-randomization test. Thus, the mean rate and its 95% highest probability density (HPD) estimated with the observed sampled dates using the Bayesian Evolutionary Analysis Sampling Trees (BEAST) v1.10.4 package (Suchard et al., 2018, see below) were compared with a null distribution obtained by randomly permutating the tip dates 10 times (Firth et al., 2010). As previously described (Duchêne et al., 2015; Murray et al., 2016), the criterion for a significant temporal signal was that the 95% HPD for the rate estimate obtained with the observed sampled dates should not overlap with the 95% HPD for the estimate obtained with randomized sampling times.

The discrete phylogeographic reconstructions were generated on the CP gene dataset using a Bayesian statistical framework implemented in BEAST v1.10.4 (Suchard et al., 2018) and the BEAGLE library (Ayres et al., 2012) to improve computational performance. BEAST uses Markov chain Monte Carlo (MCMC) integration to average over all plausible evolutionary histories for the data, as reflected by the posterior probability. The nucleotide substitution process was reconstructed using HKY85+G substitution model, the lognormal relaxed clock model and an initial substitution rate of 10^-3^ substitutions/site/year as previously determined for RYMV (Trovão et al., 2015; Dellicour et al., 2018; Issaka et al., 2021). Discrete phylogeographic inferences were estimated at the geographical level (southwestern Burkina Faso vs. the rest of West and West-Central Africa) using the continuous-time Markov chain (CTMC) process (Lemey et al., 2009) and with a Bayesian stochastic search variable selection (BSSVS). This method reconstructs the dispersal history between discrete locations and infers a posterior distribution of trees whose internal nodes are associated with an estimated ancestral location. MCMC analyses were run for one billion iterations, sampling every 100,000th and 10% iterations discarded as the chain burn-in. The maximum clade credibility (MCC) tree was obtained with TreeAnnotator v1.10.4 (BEAST package) and convergence and mixing properties (e.g., based on effective sample sizes >200 for the parameters) were inspected using Tracer v1.7.1 (http://tree.bio.ed.ac.uk/software/tracer).

### Statistical analysis of the spatio-temporal distribution of genetic groups

Contingency tables of genotype frequencies were built using the R software, and pie charts were mapped on an aerial view of the irrigated perimeter of Banzon using QGIS (QGIS 3.28, Geographic Information System; QGIS Association. http://www.qgis.org). To test whether the distribution of genotypes (frequencies of each genetic group) varied spatio-temporally (over the five sampling years and five fields), we carried out multinomial regressions with the *nnet* package (Venables & Ripley, 2002). We tested the interaction between the factor ‘year’ and the factor ‘field’ by comparing (using a likelihood ratio test in the *Anova* function of package *car;* Fox and Weisberg, 2019) the complete model (including the main effects and their interaction) with the model without the interaction, and then tested each explanatory factor.

### Phenotypic evaluation of RYMV isolates under controlled conditions

Two experiments were conducted in controlled greenhouse conditions (IRD Montpellier, France) with cycles of 12 h of light at 28°C and 80% relative humidity (RH) and 12 h of dark at 26°C and 70% RH. Five plants per pot were grown respectively in 1-L pots (Experiment 1) or 3-L pots (Experiment 2), filled with potting soil and appropriate fertilizers; soil moisture was checked throughout the experiments. RYMV isolates used in these experiments were sequenced for the CP protein (as described above), and construction of ML phylogenetic tree (see above) allowed them to be assigned to the previously identified genetic lineages.

The first experiment aimed to assess the behavior of four viral isolates, each representing one identified genetic group, on various rice cultivars. Three rice cultivars (*Oryza sativa* subsp. *indica*) commonly grown in Banzon were selected: FKR19, FKR62N and FKR64 (Barro et al., 2021a). We also included in the experiment the reference cultivars: IR64 (*O. sativa* subsp. *indica*) and Azucena (*O. sativa* subsp. *japonica*), as respectively susceptible (Albar et al., 2006) and partially resistant (Ndjiondjop et al., 1999). Three viral isolates collected at local scale in southwestern Burkina Faso from 2015 to 2019 were chosen within each of the identified genetic lineages of RYMV based on CP gene sequencing: BF706, BF710 and BF711. The isolate BF1, from the genetic lineage S2, was included as a reference severe isolate (Pinel et al., 2000).

The second experiment aimed to test the behavior of 13 viral isolates (including those used in the first experiment) from the identified genetic group on susceptible rice cultivar IR64. We used the same isolates as in experiment 1 (BF706, BF710, BF711) and also selected three additional viral isolates per genetic group. These nine isolates were sampled during other surveys conducted in Burkina Faso in 2014 (Tollenaere et al., 2017), 2016 and 2021 (unpublished): BF708, BF709, EF0738, BF712, EF0333, EF0744, BF705, EF0619 and EF0780. The BF1 isolate (Pinel et al., 2000) was also included in this experiment.

The first experiment was performed using 45 plants per rice cultivar for each of the five conditions (four viral isolates tested and the virus-free control), while we used 32 IR64 plants for each of the 14 conditions (13 viral isolates and the control) for the second experiment. In both cases, pots were arranged in two randomized blocks. The whole experiment was subdivided into three successive monthly replicates for the first experiment, while all the inoculations of the second experiment were performed at a single date.

All viral isolates were first multiplied in the susceptible rice cultivar IR64 to provide sufficient viral particles to be used as inoculum in the experiments. Then, for both experiments, viral isolates were inoculated under controlled conditions at ca. 2.10^11^ copies of viral genomes per plant. To this purpose, each inoculum was prepared by grinding infected leaves in 0.1 M phosphate buffer (pH 7.2) (Pinel-Galzi et al., 2018). Then, RYMV-specific quantitative RT-PCR (qRT-PCR) assay was used to adjust the viral loads for each inoculum, as described in Poulicard et al. (2010) with the following modifications: amplifications were carried out using 10 µL of Takyon^TM^ No Rox 2X SYBR MasterMix blue dTTP (Eurogentec, Angers, France) and performed on a LightCycler 480^TM^ Roche (Roche Diagnostics, Meylan, France). Finally, plants were mechanically inoculated 2 weeks after sowing, using 50 µL of inoculum per leaves for each RYMV isolates, or inoculated with phosphate buffer alone (virus-free control), as described in Pinel-Galzi et al. (2018).

Infected plants were monitored for visual symptoms beginning at 9 days post-inoculation (dpi) and every 3-4 days until 24 dpi. Yellow mottle symptoms appeared on newly developed leaves after 10 days at the earliest and were scored according the symptom severity scale described in Pinel-Galzi et al. (2018). In order to estimate the intra-plant viral load for each condition, early in the infection process, we sampled meristem from one infected plant per pot at 7 dpi and performed a qRT-PCR assay as described previously. The number of RYMV copies was estimated using a standard curve obtained with purified virus (BF5 isolate, S1wa strain).

### Experimental infection data analysis

The overall aggressiveness of viral isolates on plants was evaluated by considering the severity of symptoms and viral load. For each inoculated plant, symptoms at 9, 13, 16, 21 and 24 dpi were used to compute the area under the disease progression curve (AUDPC) using the R package *agricolae* (De Mendiburu and Yaseen, 2020).

Results obtained for each of the three replicates of the first experiment were analyzed jointly for statistical analysis of AUDPC and viral load. The R package *ggplot2* was used to visualize the effect of the virus on symptom expression (AUDPC) and viral load (of RYMV copy number at 7 dpi expressed in logarithm) for each RYMV isolates on the different rice cultivars.

Both experiments were analyzed with GLMM using R package *lme4*. For the first experiment, the effects of the variables ‘RYMV isolates’, ‘rice cultivars’ and their interaction, were tested independently on the AUDPC and the viral load (number of RYMV copies at 7 dpi). Inoculation date was included in the models as random factor. For the second experiment, we tested the effect of the variables ‘RYMV isolates’ and ‘genetic group’ on the variables AUDPC and viral load, including the variable ‘block’ as a random factor. Post hoc tests were performed using the R package *lsmeans*.

## RESULTS

### Yellow mottle disease prevalence varies spatio-temporally in western Burkina Faso

A total of 203 observations were performed between 2015 and 2019, with total number of studied field per year varying between 24 and 50 (Table 1). The total number of fields presenting yellow mottle disease symptoms was 62, representing a frequency of 30.5% over the whole area and period. In each studied site and year, the frequency of fields where yellow mottle disease symptoms were found varied from 0 to 87.5% (Table 1), with a maximum of 7 fields with symptoms out of 8 studied fields in Banzon in 2017. Detailed results obtained for yellow mottle disease symptoms, observed in the four cells of the diagonal of the grid on the one hand, and in the 16 plants of grid nodes on the other hand, are available on DataSuds https://doi.org/10.23708/GZCM1O.

**Table 1:** Frequency of studied fields presenting yellow mottle disease symptoms and median and average prevalence in the six sites studied from 2015 to 2019 in western Burkina Faso. The number of fields were yellow mottle disease were observed are mentioned over the total number of rice fields were symptom observations were performed are indicated for each year. Median and average prevalences are based on RYMV-specific DAS-ELISA test in all studied fields (including both fields with and without yellow mottle disease symptoms observed). The different sites are grouped by geographical zones, see Figure 1. RL: Rainfed Lowland; IR: irrigated areas.

| Site | Number of fields with yellow mottle disease symptoms observed (%) |  |  |  |  |  |
| --- | --- | --- | --- | --- | --- | --- |
|  | Median and average prevalences based on DAS-ELISA results |  |  |  |  |  |
|  | 2015 | 2016 | 2017 | 2018 | 2019 | All years (2015-19) |
| Bama-RL<br>(Badala) | NA | 4/6 (66.7%)<br>0% - 2.1% | 4/8 (50.0%)<br>0% - 13.3% | 2/7 (28.6%)<br>0% - 1.8% | 3/6 (50.0%)<br>0% - 2.1% | 13/27 (48.1%)<br>0% - 5.3% |
| Bama-IR<br>(Bama) | NA | 0/10 (0%)<br>0% - 0% | 1/8 (12.5%)<br>0% - 4.7% | 1/9 (11.1%)<br>0% - 0.7% | 0/7 (0%)<br>0% - 0% | 2/34 (5.9%)<br>0% - 1.3% |
| Banzon-RL<br>(Senzon) | NA | 1/5 (20.0%)<br>0% - 1.3% | 2/7 (28.6%)<br>0% - 1.8% | 1/7 (14.3%)<br>0% - 0.9% | 2/7 (28.6%)<br>0% - 2.7% | 6/26 (23.1)<br>0% - 1.7% |
| Banzon-IR<br>(Banzon) | 4/12 (58.3%)<br>0% - 18,2% | 6/8 (75.0%)<br>3.1% - 9.4% | 7/8 (87.5%)<br>71.9% - 54.7% | 9/11 (81.8%)<br>12.5% - 17.6% | 3/6 (50%)<br>9.4% - 17.7% | 32/45 (71.1%)<br>12.5% - 22.9% |
| Karfiguela-RL<br>(Tengrela) | NA | 1/5 (20.0%)<br>0% - 0% | 2/7 (28.6%)<br>0% - 2.7% | 1/7 (14.3%)<br>0% - 8.0% | 2/6 (33.3%)<br>0% - 13.5% | 6/25 (24.0%)<br>0% - 6.25% |
| Karfiguela-IR<br>(Karfiguela) | 1/12 (8.3%)<br>0% - 0% | 1/9 (11.1%)<br>0% - 0.7% | 1/8 (12.5%)<br>0% - 0% | 0/9 (0%)<br>0% - 0% | 0/8 (0%)<br>0% - 0% | 3/46 (6.5%)<br>0% - 0.1% |
| Total<br>(all sites) | 8/24 (33.3%)<br>0% - 9.1% | 13/43 (30.2%)<br>0% - 2.3% | 17/46 (36.9%)<br>0% - 13.3% | 14/50 (28%)<br>0% - 5.5% | 10/40 (25%)<br>0% - 5.5% | 62/203 (30.5%)<br>0% - 7.0% |

Serological analysis was performed for the 16 plants collected in the 62 fields where yellow mottle disease symptoms were observed, resulting in a dataset of 990 samples (62x16 and 2 missing data). We observed a total of 228 positive plants (global seroprevalence: 23.0%) when applying a constant threshold (OD = 0.3) (see also N’Guessan et al., 2000), while 353 plants (global seroprevalence: 35.7%) were DAS-ELISA-positive when applying a threshold following (Traoré et al., 2008). Comparing serological results with the presence/absence of observed symptoms led us to choose the constant threshold for subsequent data analysis. Indeed, with the approach of Traoré et al. (2008), we obtained 107 DAS-ELISA-positive plant where no symptom could be observed (30.3% of DAS-ELISA-positive plants) and 16 negative symptomatic plants (1.6% of total analyzed plants). On the other hand, with a constant threshold, we obtained only 28 ELISA-positive plants where no symptom could be observed (12.3% of DAS-ELISA-positive plants) and 20 negative symptomatic plants (2.0% of total analyzed plants). Statistical analyses were however performed also with the approach of Traoré et al. (2008) and the conclusions did not change (data not shown).

In the fields where we did not observe any yellow mottle disease symptom, we assumed the disease was absent and set prevalence to zero. The distribution of RYMV prevalence based on DAS-ELISA results over the six studied sites and the five-year period appears in Table 1 and Figure 1. The relationship between prevalence estimates based on DAS-ELISA data and symptom-based prevalence data is shown in Supplementary Figure 1. It revealed highly significant correlations, both when symptoms are observed on the same 16 plants (cor = 0.97; *p* < 0.001) and when symptoms are observed in the four diagonal grid cells (cor = 0.78; *p* < 0.001). The interaction between rice production system and geographical zone, that reflect the particular site (among six sampled sites), had a highly significant effect (*p* < 0.0001) on detection result, as well as the factor ‘Year’, both including or not the fields with no yellow mottle disease symptoms at all (see Supplementary Table 1).

**Figure 1:**
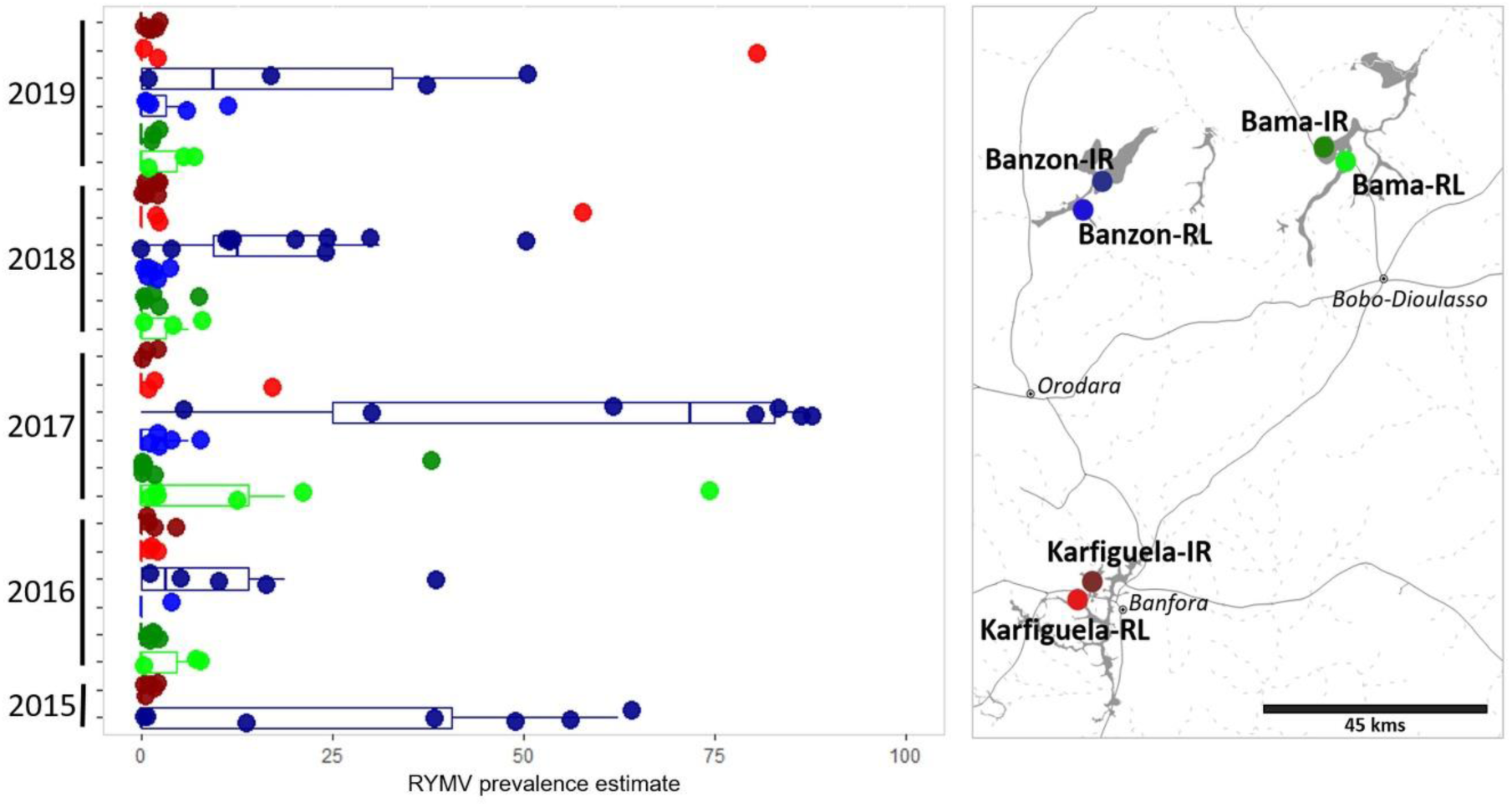
Rice yellow motte virus (RYMV) prevalence data over the different sites in western Burkina Faso within the 2015-2019 period. RYMV prevalence estimate are derived from specific serological detection of RYMV in 16 plants analyzed in each field where yellow mottle disease symptoms were observed.

For all five studied years, the site presenting the highest RYMV prevalence (significantly different from all other five sites) is the irrigated perimeter of Banzon (Banzon-IR, Figure 1 and Supplementary Figure 2). In Banzon-IR site, 36.9% of studied fields presented yellow mottle symptoms, with average prevalence up to 13.3%, over the 2015-2019 period (Table 1). Also, Bama-RL and Karfiguela-RL had quite high frequency of infected fields and average prevalences, while the opposite was found for the irrigated perimeters of Bama and Karfiguela (Bama-IR and Karfiguela-IR, Table 1).

In terms of temporal variation, between-year differences were all significant, except when comparing 2018 and 2019. We observe the highest frequency of infected fields (36.9%) and average prevalence (13.3%) in 2017, while the lowest average prevalence (2.3%) was observed in 2016 (see Table 1 and Supplementary Figure 2).

### Agricultural practices affect yellow mottle disease prevalence in a disease hotspot

GLMM performed to explain the rice yellow mottle virus (RYMV) prevalence within the irrigated perimeter of Banzon revealed an effect of the rice cultivar (χ² = 14.3, *p* = 0.002), but no effect of the off-season cultivation (χ² = 0.15, *p* = 0.69), while a significant effect of their interaction is observed (χ² = 7.86, *p* = 0.049). Tukey post hoc tests for the variable ‘rice cultivar’ revealed that FKR64 differ from FKR34 (Z = 2.77, *p* = 0.025) and FKR62N (Z = -2.89, *p* = 0.018); and FKR34 and FKR62N differ from each other (Z = 3.74, *p* < 0.001). Lower RYMV prevalences are observed in fields known as cultivated with FKR64, while higher RYMV prevalences are found for FKR34 (Supplementary Figure 3). In addition, both the double rice season (compared to only one rice cultivation per year) and various fertilization applications (two or more, as compared to one or less) correspond to higher RYMV prevalence (Supplementary Figure 3).

### Yellow mottle disease aggregation is observed within small rice fields

Within the small (20×20 m) studied fields, there were 15% more pairs of infected plants in the first distance class (5 m) than expected under the null hypothesis of random distribution of the infection, both for observed yellow mottle disease symptoms (*p* < 0.001) and for serological RYMV detection (*p* < 0.001). This pattern of aggregation between infected plants remains significant (*p* < 0.05) even within a 10-m radius.

### The disease hotspot harbors high genetic diversity

A total of 132 CP gene sequences (clear chromatograms) were generated from samples collected in Banzon between 2015 and 2019, with number of samples per field and year varying between 0 and 16 (average: 5.3±4.3; Table 2). In addition, 6 samples (4.3% of the dataset) were amplified but resulted in unclear chromatogram, possibly as a result of mixed infection by various isolates.

**Table 2:** List of the five fields studied from 2015 to 2019 within the irrigated area of Banzon with the estimated RYMV prevalence and number of plants for which CP sequence was obtained. RYMV prevalences are based on RYMV-specific serological test (DAS-ELISA). The last line (Total, all fields) refers to all studied fields in the irrigated area of Banzon (not only the five fields shown here).

| Field | RYMV prevalences |  |  |  |  |  |
| --- | --- | --- | --- | --- | --- | --- |
|  | Number of plant samples with CP sequence in each field and year |  |  |  |  |  |
|  | 2015 | 2016 | 2017 | 2018 | 2019 | Total<br>(average all<br>years) |
| BZ02 | 62.5% | 6.2% | 0% | 12.5% | 0% | 16.2% |
|  | 5 | 3 | 0 | 4 | 0 | 12 |
| BZ04 | 12.5% | 0% | 87.5% | 25.0% | 50.0% | 35.0% |
|  | 4 | 0 | 6 | 4 | 16 | 30 |
| BZ09 | 37.5% | 37.5% | 87.5% | 50.0% | 0% | 42.5% |
|  | 6 | 5 | 5 | 9 | 0 | 25 |
| BZ10 | 56.2% | 18.7% | 81.2% | 12.5% | 18.7% | 37.5% |
|  | 10 | 3 | 4 | 10 | 10 | 37 |
| BZ11 | 50.0% | 0% | 81.2% | 6.2% | 37.5% | 35.0% |
|  | 4 | 4 | 4 | 1 | 15 | 28 |
| Total (all<br>fields) | 18.2% | 9.4% | 54.7% | 17.6% | 17.7% | 22.9% |
|  | 29 | 15 | 19 | 28 | 41 | 132 |

Interestingly, the genetic diversity of the 132 CP gene sequences observed in Banzon (d=0.0391±0.0051 nucleotide substitutions per site) is similar to those estimated for the whole Burkina Faso (d=0.0449±0.0054 sub/site) and is higher than those from most countries in West- and West-Central Africa (Supplementary Figure 4), which could reflect a complex or old epidemiological history of RYMV in this region.

### Various historical introductions shaped RYMV viral diversity in southwestern regions of Burkina Faso

Based on the CP dataset collected in Banzon (132 CP) and in West- and West-Central Africa (261CP, including some sequences from several regions of Burkina Faso), we reconstructed the evolutionary history of RYMV to gain information on the epidemic history of this virus in southwestern Burkina Faso. First, we established that no recombination event (*p* = 0.93) was detected within the CP dataset (393 sequences in total). Then, the strength of the temporal signal was examined on this CP dataset by linear regression exploration of root-to-tip distances and by date-permutation tests. The linear regression exploration showed a low but significant temporal signal in the 393 CP dataset (*R*² = 0.087; *p* = 1.62 x 10^-9^; Supplementary Figure 5). The presence of temporal signal was confirmed with the date-randomization tests (tip-date randomization: Supplementary Figure 6A and B; clustered tip randomizations: Supplementary Figure 6C and D) using BEAST, as none of the estimates of the permutations overlapped with the ones of the real (non-permutated) 393 CP dataset (Supplementary Figure 6).

We reconstructed the discrete dispersal histories of the RYMV in West- and West-Central Africa by Bayesian inferences. We obtained a maximum clade credibility (MCC) tree where the tip locations (southwestern Burkina Faso vs. the rest of West- and West-Central Africa) were mapped (Figure 2A). Eight independent introductions of RYMV to southwestern Burkina Faso were estimated (*MarkovJumps*_Introduction_ = 8.11 [8.00; 9.00]; branches 1 to 8 indicated in Figure 2A), two corresponding to S2 strain introductions (branches 1 and 2), three to S1wa strains (branches 3 to 5) and three to S1ca strains (branches 6 to 8). We also distinguish branch 9 that corresponds to a group named S1bzn. The date of emergence of each RYMV strain in southwestern Burkina Faso could be estimated based on the date-intervals of the nodes of branches of interest. The introductions of S2 and S1wa strains in this region were estimated between the late 1980’s to the 2000’s/2010’s (Figure 2B, Table 3). The introduction of the S1ca strain probably occurred later, from the 2000’s to mid 2010’s (Figure 2B, Table 3). Finally, the emergence of the group S1bzn was estimated to the beginning of the 2010’s (Figure 2B, Table 3).

**Figure 2:**
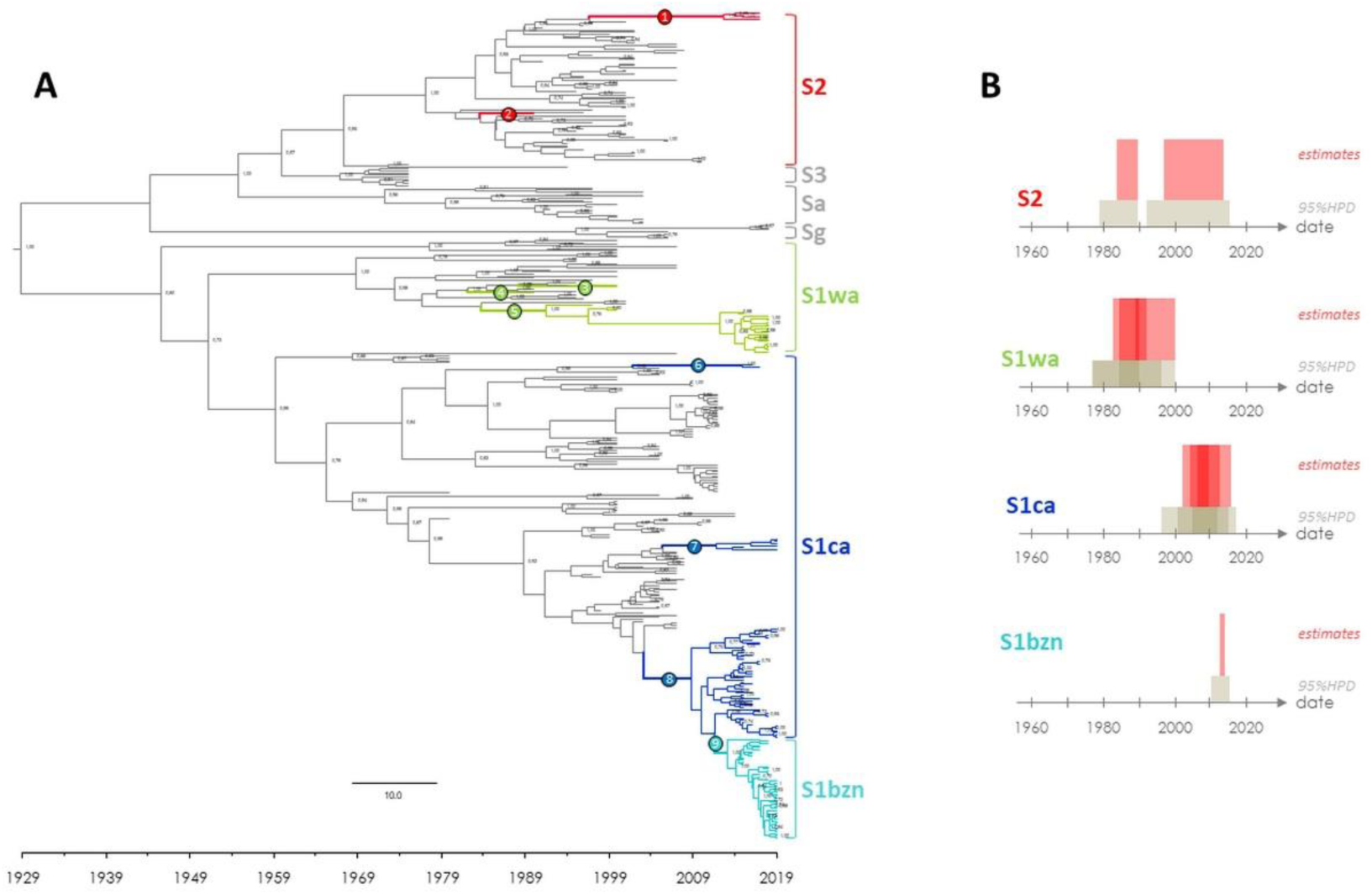
(A) Time-calibrated maximum clade credibility (MCC) tree reconstructed by continuous Bayesian evolutionary inference with the 393 coat protein (CP) gene sequence dataset. Branches corresponding to the isolates collected in Southwestern Burkina Faso are colored according to the RYMV strains (S1wa: green; S1ca: blue; S1wa x S1ca recombinant line named S1bzn: light blue; S2: red). The values on the nodes correspond to their posterior probabilities (values above 0.70 are indicated). The bottom axis gives the timeframe (by date) of the RYMV diversification. Nine branches of interest are labeled on the MCC tree (from 1 to 9) and their relative nodes (1a, 1b, 2, 3, 4, 5a, 5b, 6a, 6b, 7a, 7b, 8a, 8b, 9a, 9b) were annotated with their time to the most common ancestor and their 95% highest probability density (HPD) intervals (Table 3). (B). Dates (in red) and the 95% highest probability densities (HPD) intervals (in grey) of the introduction/emergence of the S2, S1wa, S1ca and recombinant line S1bzn in Southwestern Burkina Faso estimated as the intervals between nodes of interest (Table 3). The darker areas represent the overlaps between the estimates.

**Table 3:** Introduction dates of RYMV strains in Southwestern Burkina Faso. Date-intervals for nine branches of interest and their relative nodes (1a, 1b, 2, 3, 4, 5a, 5b, 6a, 6b, 7a, 7b, 8a, 8b, 9a, 9b), annotated with their time to the most common ancestor and their 95% highest probability density (HPD) intervals.

| *Time before present (2019,83)* |  |  |  |  |  |
| --- | --- | --- | --- | --- | --- |
| N° node | Time | Time 95%HPD | Date | Date 95%HPD | Genetic group |
| **1a** | 22,71 | [27.79 - 18.18] | **1997,12** | [1992.04 - 2011.65] | **S2** |
| **1b** | 6,59 | [9.05 - 4.53] | **2013,24** | [2010.78 - 2015.30] |  |
| **2** | 35,86 | [40.97 - 31.29] | **1983,97** | [1978.86 - 1988.54] |  |
| **3** | 30,94 | [35.24 - 26.59] | **1988,89** | [1984.59 - 1993.24] | **S1wa** |
| **4** | 37,04 | [42.40 - 32.51] | **1982,79** | [1977.43 - 1987.32] |  |
| **5a** | 35,32 | [43.07 - 27.82] | **1984,51** | [1976.76 - 1992.01] |  |
| **5b** | 27,88 | [33.22 - 23.48] | **1991,95** | [1986.61 - 1996.35] |  |
| **6a** | 17,90 | [23.34 - 14.29] | **2001,93** | [1996.49 - 2005.54] | **S1ca** |
| **6b** | 4,48 | [6.85 - 3.05] | **2015,35** | [2012.98 - 2016.78] |  |
| **7a** | 13,74 | [15.02 - 12.92] | **2006,09** | [2004.81 - 2006.91] |  |
| **7b** | 7,53 | [10.07 - 5.13] | **2012,30** | [2009.76 - 2014.70] |  |
| **8a** | 15,77 | [18.97 - 13.43] | **2004,06** | [2000.86 - 2006.40] |  |
| **8b** | 10,65 | [13.80 - 8.23] | **2009,18** | [2006.03 - 2011.60] |  |
| **9a** | 7,56 | [9.64 - 6.07] | **2012,27** | [2010.19 - 2013.76] | **S1bzn** |
| **9b** | 5,91 | [7.39 - 4.79] | **2013,92** | [2012.44 - 2015.04] |  |

Note that no migration of RYMV populations from southwestern Burkina Faso to other regions of West- and West-Central Africa was reported using this dataset (*MarkovJumps*Release = 0.11 [0.00; 1.00]).

### Co-circulation of viral genetic groups, with temporal variation throughout the sampling period

The minimum spanning network (MSN) connecting all the CP sequences (at the amino-acid level) from Banzon is presented in Figure 3. While the genetic differentiation is low between fields (*F*_ST(field)_ = 0.0562; *p* = 0.0134) and rice cultivars (*F*_ST(rice)_ = 0.0474; *p* = 0.0098), it is highly significant between years (*F*_ST(years)_ = 0.2796; *p* < 0.0001, Figure 2). Notably, the genomes identified in 2018 and 2019 were significantly different from the genomes sampled the other years, suggesting wide variations on RYMV populations circulating in Banzon during the last two sampling years.

**Figure 3:**
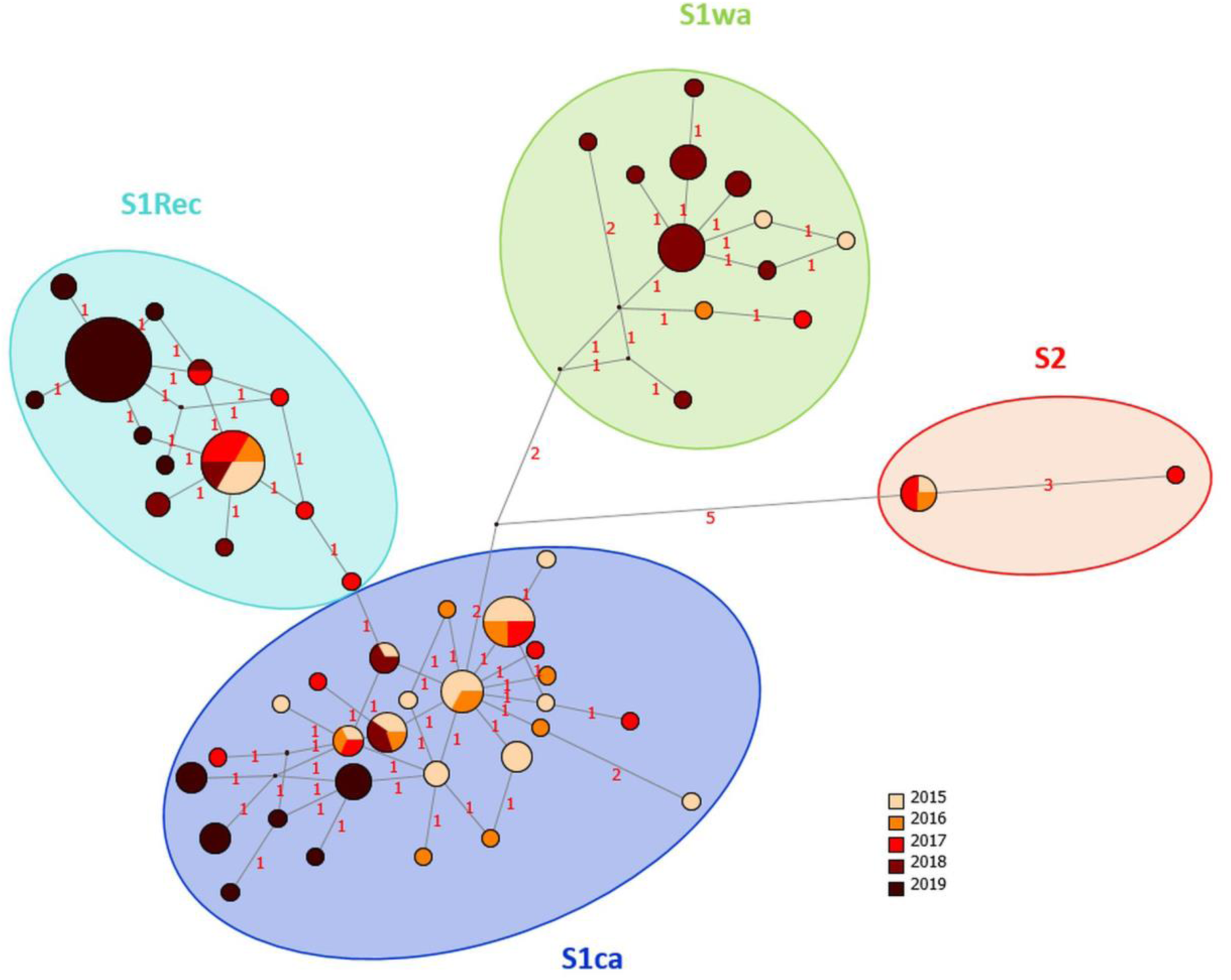
Minimum spanning network (MSN) of the 132 coat protein (CP) sequences (at the amino acid level) identified in Banzon. Each CP sequence is indicated by one node in which the circle area is proportional to the number of individual sequences for this particular sequence and the year of sampling of these sequences is represented as a proportional pie chart. The number amino acid substitutions between sequences are indicated on the branches. Seven putative intermediate sequences connecting the CP sequences correspond to the small red dots.

### Three genetic groups are distinguished on the network (Figure 3) and named based on the phylogenetic tree including reference isolates from West- and West-Central Africa (Figure 2A): S1ca, S1wa and S2

A fourth group, corresponding to lineage 9 in the phylogenetic tree, is labeled S1bzn. This group presents a clear temporal structure with genotypes that progressively diverge from the rest of the network through time (Figure 3).

Whole genome sequencing was performed for one isolate from each of the three major genetic groups (S1wa, S1ca, S1bzn), and the three newly obtained complete sequences were compared to the 37 RYMV genomes described from samples collected in West- and West-Central Africa (Supplementary Figure 7). For the samples of group S1bzn (2017MP1493), the software RDP identified one putative recombination event, with a CP from the S1ca group while the rest of the genome corresponds to the S1wa group. In accordance with this detected recombination event, we obtained incongruent phylogenies for the CP sequences and for the rest of the genome (Supplementary Figure 7).

Contingency tables of the four genetic groups are presented in the Supplementary Table 2, and pie charts are represented in each field and year (Figure 4). Various genetic groups were found within the same field, with up to three genetic groups co-existing at the same sampling date (Figure 4 and Supplementary Table 2). Multinomial regressions showed no effect of the interaction between year and field, and no geographical structure within the irrigated perimeter (factor ‘field’ not significant). On the other hand, a strong effect of the factor ‘year’ is evidenced (χ² = 95.2; df =12; *p* < 0.001). Temporal variations in the frequency of genetic groups are presented in Figure 5. The overall frequency over the 5 years are: 44.6% for the genetic group S1ca, 37.1% for S1bzn, 15.2% for S1wa and only 3.0% for S2. The three groups S1wa, S1ca and S1bzn coexist throughout the five-year period with fluctuations in time, while the S2 strain was only found in 2015 and 2017. The S1wa strain was rare in 2015 and absent in 2016, 2017 and 2019; however, it was the most frequent lineage in 2018 (64%). A decrease in S1ca frequency is observed over the period (from 76% in 2015 to 32% in 2019), whereas S1bzn was rare in 2015 and 2016 (14-20%) but seemed to rise in 2019 (up to 68%). Finally, the overall (2015-2019) genetic diversity in the studied site is 0.039 ± 0.005 sub/site, and varies in time with a maximum reached in 2017 (0.047 ± 0.005 sub/site) and 2018 (0.042 ± 0.005 sub/site) and a minimum in 2019 (0.015 ± 0.003 sub/site; Figure 5).

**Figure 4:**
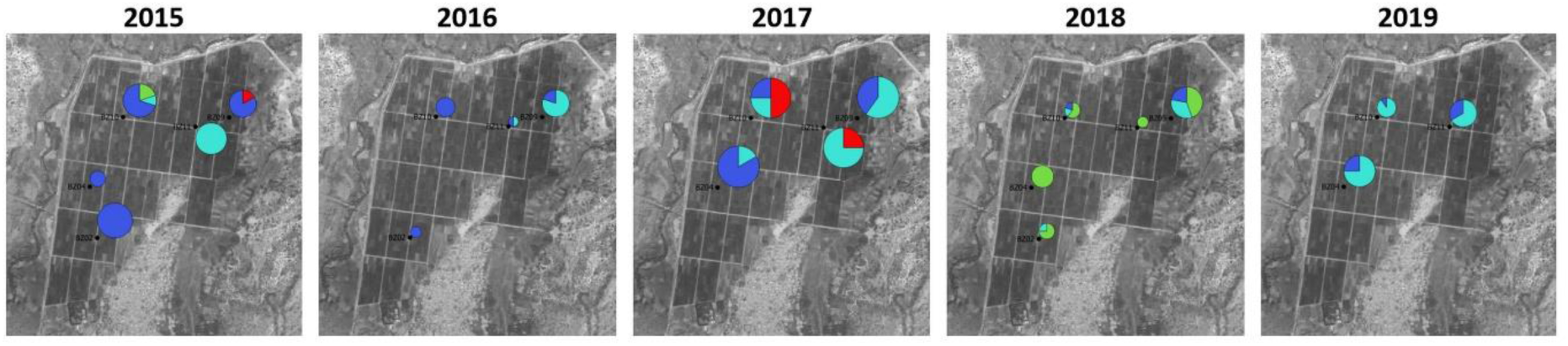
RYMV prevalence and genetic diversity (CP sequences) follow-up data over the five selected quadrats within the irrigated area of Banzon between 2015 and 2019. The colors represent the four identified genetic groups: S2 in red, S1wa in green, S1ca in dark blue and the recombinant S1bzn in light blue (see also Supplementary Table 2). The surface of each pie chart represent RYMV prevalence estimate based on serological data (see also Table 2). The only exception is for BZ11 in 2016, where not any of the 16 plants analyzed where positive in DAS-ELISA, while a few symptomatic plants were observed and sequenced (Table 2), and we set the prevalence to 5 in this field to be able to represent the genetic groups identified.

**Figure 5:**
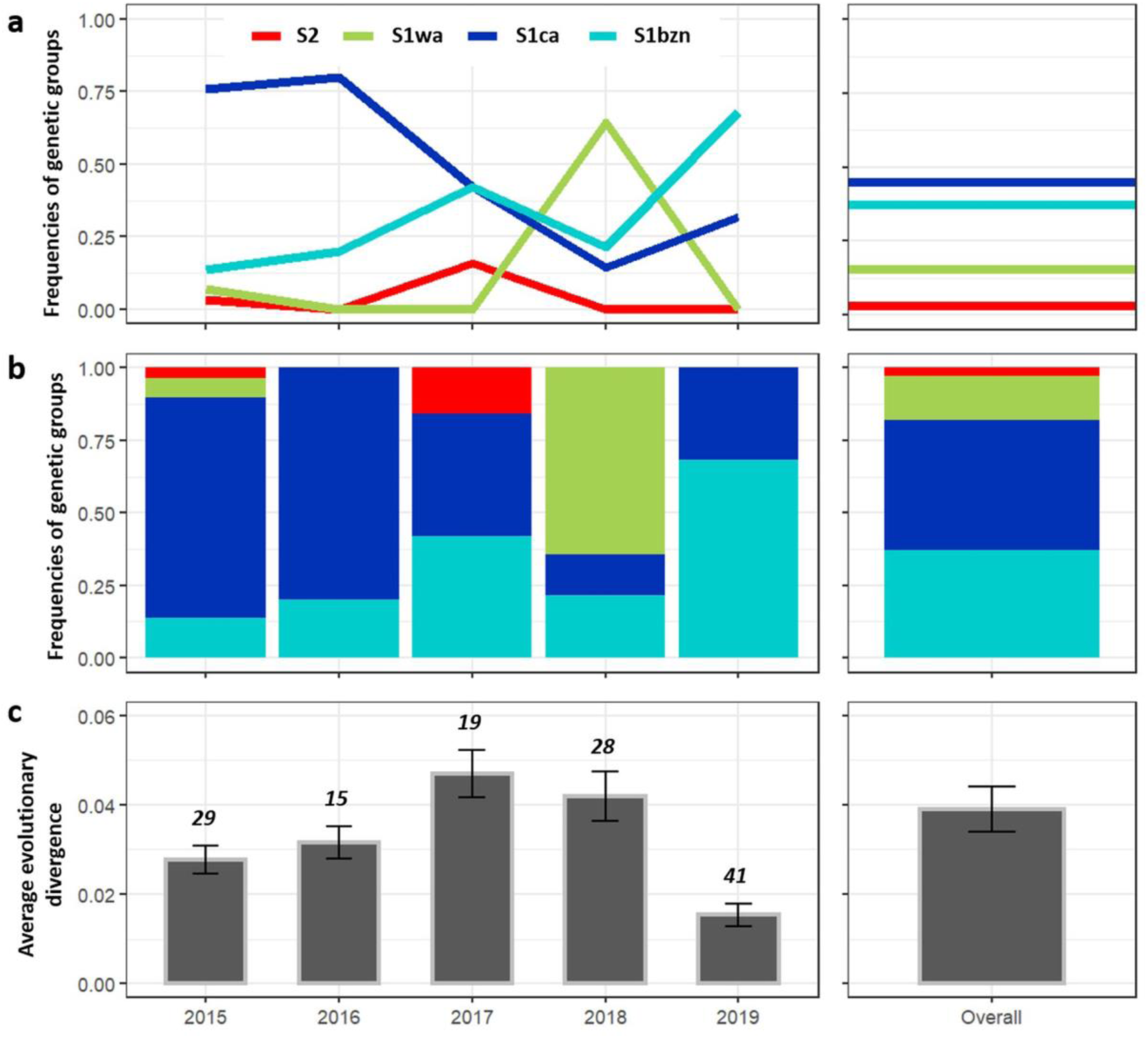
Temporal variation of RYMV genetic diversity (coat protein region) over the 2015-2019 period in the irrigated perimeter of Banzon. a and b. Overall frequencies of the four genetic groups in Banzon for the year 2015, 2017, 2018 and 2019. The four genetic group are represented by four different colors with S2 in red, S1wa in green, S1ca in dark blue, and S1bzn in light blue. c. Estimates of Average evolutionary divergence over sequence pairs within groups. The number of base substitutions per site from averaging over all sequence pairs within each group are shown. Standard error estimate(s) are shown above the diagonal. Analyses were conducted using the Kimura 2-parameter model. The right panel labelled ‘Overall’ represents the whole dataset of 132 CP sequences sampled over the 2015-2019 period.

### Symptoms and viral load are affected by the rice cultivar, the viral isolate and their interactions

Detailed results obtained for all experimental infections performed are available on DataSuds preliminary link: https://doi.org/10.23708/MDQIUE. The phylogenetic position of the different isolates used in experiments are presented in Supplementary Figure 8 and Supplementary Table 3: isolates BF706, EF0738, BF708 and BF709 belong to the S1wa strain. BF710, EF0744, BF712, EF0333 are identified as S1ca and EF0619, EF0780, BF705, BF711 as S1bzn.

Firstly, in order to characterize the pathogenicity of the major viral genetic groups identified, we evaluated the symptom severity and intra-plant viral accumulation of four RYMV isolates from each genetic group identified (S2, S1wa, S1ca, S1bzn) on the rice cultivars commonly grown in Burkina Faso (FKR19, FKR62N, FKR64) and 2 control varieties (susceptible rice cultivar IR64 and partially resistant cultivar Azucena) (Supplementary Figure 9).

GLMM performed to explain the symptom severity on the one hand and viral load on the other hand revealed in both cases an effect of the rice cultivar, the RYMV isolate, and the interaction between these two factors (Table 4). Tukey post hoc tests revealed that IR64 and Azucena differ from other rice cultivars for both symptom severity and viral load (respectively *p* < 0.0001 and *p* < 0.05; Supplementary Table 4). Highest symptoms and viral load are observed for the four RYMV isolates inoculated on IR64, while the lowest symptoms and viral load are observed on the rice cultivar Azucena. On the opposite, a common intermediate pattern is observed for the three cultivars from Burkina Faso (FKR19, FKR62N and FKR64), both for symptoms and viral load. Indeed, Tukey post hoc tests revealed no significant differences between the three rice cultivars for symptoms and viral load (Supplementary Table 4).

**Table 4:** Statistical analyses of the experimental infection results: estimations of symptom severity and viral load in a set of 4 isolates and 5 cultivars (Experiment 1) and in a set of 13 isolates on rice cultivar IR64 (Experiment 2). Results of the GLMM for the Experiment 1 including ‘Inoculation date’, ‘Block’, ‘Rice cultivar*Isolate’ and for the Experiment 2 including ‘Block’ and ‘Genetic group/Isolate’ are reported. GLMM results on symptom severity (integrated over the symptom development: AUDPC = Area Under the Disease Progress Curve) are shown on the left, and on viral load (estimated from specific qPCR on meristematic area at 7dpi) on the right, for each experiment.

Experiment 1
| Model (GLMM) | AUDPC |  |  | Viral load |  |  |
| --- | --- | --- | --- | --- | --- | --- |
|  | Chisq | Df | Pr(>Chisq) | Chisq | Df | Pr(>Chisq) |
| Rice cultivar | 566.957 | 4 | <2.000e-16 | 141.486 | 4 | <2.200e-16 |
| Isolate | 81.238 | 3 | <2.000e-16 | 64.870 | 3 | 5.349e-14 |
| Rice cultivar:Isolate | 23.728 | 12 | 0.022 | 30.643 | 12 | 0.002 |

Experiment 2
| Model (GLMM) | AUDPC |  |  | Viral load |  |  |
| --- | --- | --- | --- | --- | --- | --- |
|  | Chisq | Df | Pr(>Chisq) | Chisq | Df | Pr(>Chisq) |
| Genetic group | 1.256 | 3 | 0.740 | 5.813 | 3 | 0.121 |
| Genetic group:Isolate | 1146.548 | 9 | <2.000e-16 | 61.704 | 9 | 6.285e-10 |

For the five rice cultivars tested, we observed similar pattern of variability in aggressiveness according to RYMV isolates (Supplementary Figure 9). Firstly, the BF1 isolate (S2 strain) is the most aggressive: it caused the strongest symptoms and accumulated to the highest level in all rice cultivars. In contrast, isolate BF711 (from lineage S1bzn) is the least aggressive isolate, with mildest symptoms and low level of accumulation in all rice cultivars. Tukey HSD tests, all presented in Supplementary Table 4, showed highly significant differences between BF1 and BF711, both for symptoms (t = 8.44, *p* < 0.0001) and viral load (t = 7.99, *p* < 0.001). Intermediate behavior was observed for isolate BF706 (S1wa strain), which is different from BF1 (S2) and BF711 (S1bzn) for both symptoms and viral load (see Tukey HSD tests; Supplementary Figure 4). Finally, the BF710 isolate (S1ca strain), caused symptoms with no differences from BF1 (S2) and BF706 (S1wa). An intermediate level of accumulation in all rice cultivars was observed for BF710 (S1ca) which is not different from BF706 (S1wa) but differs from BF1 (Tukey HSD, *p* < 0.0001; Supplementary Figure 4).

### Isolates from S1bzn group induced severe symptoms and high viral load

A second experiment was performed to evaluate potential variability in pathogenicity within genetic groups. To this purpose, we evaluated on the susceptible rice cultivar IR64 the experimental behavior of three additional isolates for each of three genetic groups (S1wa, S1ca and S1bzn). GLMM results revealed that the genetic group was not a determinant of either symptom severity or viral load; only the specific RYMV isolate has an effect on symptom severity and viral load (Table 4). The Figure 6 shows the ranking of all tested isolates, in terms of aggressiveness and viral accumulation (see also the contrast analysis in Supplementary Table 4).

**Figure 6:**
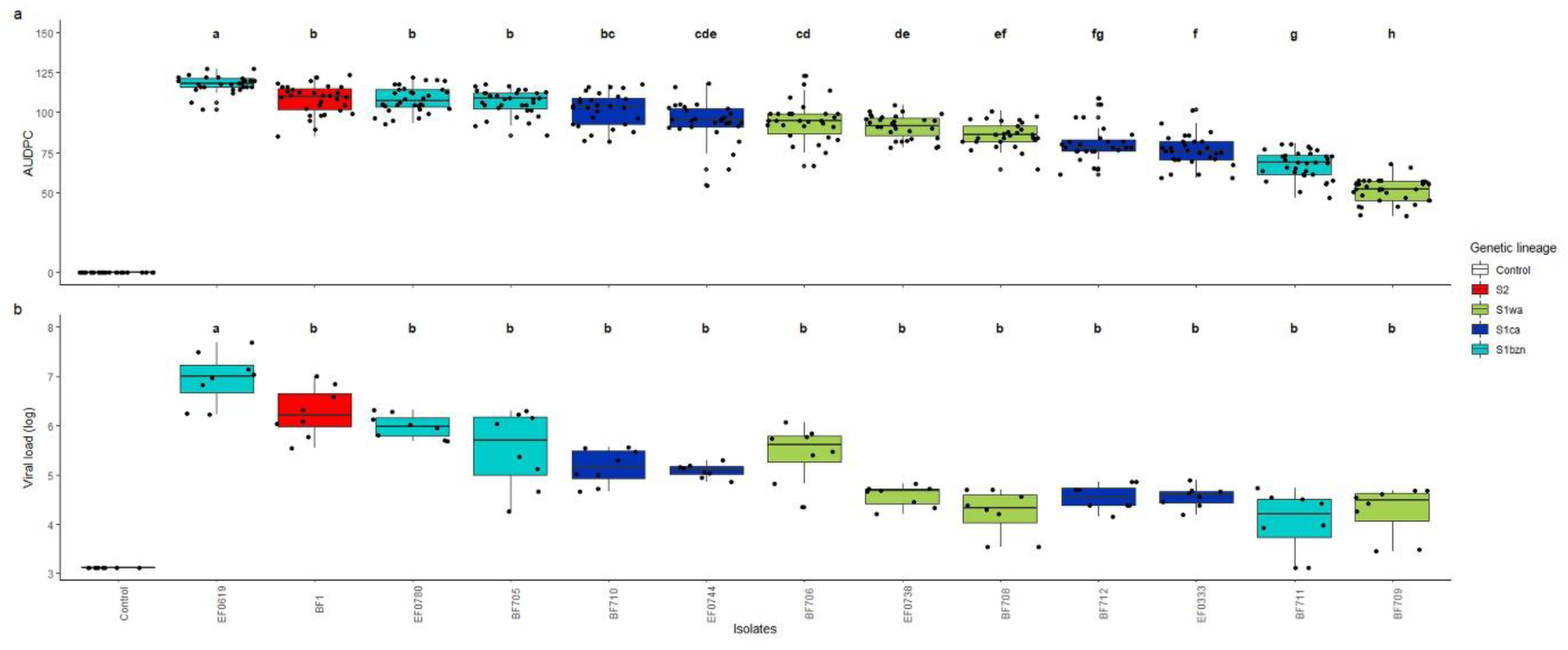
Symptom estimate and viral load of 13 RYMV isolates, from the three genetic groups identified in Banzon (S1wa, S1ca, S1bzn), evaluated on a susceptible rice cultivar. a. The symptom severity estimated by the area under disease progress curve (AUDPC) of 13 RYMV isolates inoculated on the rice cultivar IR64. AUDPC was determine according the notation severity scale over five dates (from 9 to 24 dpi), on 32 phenotyped plants. b.The intra-plant viral load of the 13 RYMV isolates estimated on the rice cultivar IR64. The number of RYMV copies was estimated by RT-qPCR on 8 meristematic area collected at 7 dpi, using a standard curve. Isolates are ordered according to their aggressiveness (highest to lowest AUDPC value). The 13 RYMV isolates are represented by four different colors corresponding to their genetic group, S2 (red), S1wa (green), S1ca (dark blue) and S1bzn (light blue). Letters indicate the significant differences between each isolate (Tukey honestly significant difference, *p* < 0.05).

For the four isolates tested in the first experiment, we obtained the same pattern of infection outcome as in Experiment 1: BF1 (S2) being the most aggressive and BF711 (S1bzn) the least aggressive. In addition, this second experiment shows that the isolate causing the highest symptoms and accumulating at the highest level in IR64 is EF0619 from S1bzn group (Tukey HSD, *p* < 0.001; Supplementary Table 4). The genetic group S1bzn also contains two other aggressive isolates (EF0780, BF705), so that out of four isolates tested from this group, three induce strong symptoms and accumulate at high levels.

## DISCUSSION

Pathogens offer the unique opportunity to study contemporaneous evolution, and monitor “evolution-in-action” (McDonald, 2010), a fascinating context for evolutionary biology. Understanding pathogen evolution is also a priority for public health, as well as food security, especially for virus diseases affecting subsistence agriculture in developing countries (Jones & Naidu, 2019). In this context, we document here the spatio-temporal variation in prevalence, as well as genetic and pathogenic diversity, of the rice yellow mottle disease at local scale in western Burkina Faso.

Serological detection of RYMV performed in all the studied fields where yellow mottle disease symptoms were observed, in a survey of six sites over five years, confirmed previous results based on symptoms: the rice growing system is not a structuring factor for the rice yellow mottle disease in western Burkina Faso (Barro et al., 2021a). Indeed, the irrigated perimeter of Banzon is identified as the most frequently affected by the disease in this region, and where prevalence is higher. It confirms this site as a hotspot of yellow mottle disease as previously noticed (Traoré et al., 2015; Barro et al., 2021a). In addition, two rainfed lowland sites (Bama-RL and Karfiguela-RL) occasionally experience quite high frequency of infected fields and prevalences, but much less than Banzon-IR. This geographical pattern of yellow mottle disease is consistent with farmers interviews data. Indeed, rice diseases were mentioned among the major threats to rice production in 12 cases out of 83 (14.5%) over the six study sites, 7 of which (58%) were from the irrigated perimeter of Banzon. Among the diseases known by the farmers, yellow mottle disease was cited in 25 cases out of 82 (30.5%), 18 (72%) of which were from the irrigated perimeter of Banzon. Consequently, these results confirm that the site where disease burden is highest corresponds to the site where the farmers recognize yellow mottle disease, as also shown in a previous study of RYMV in Burkina Faso (Traoré et al., 2015). The disease control methods mentioned by the farmers in the irrigated perimeter of Banzon are: (i) uprooting of symptomatic plants, (ii) avoiding entering a rice field if symptoms are observed to avoid further plant-to-plant transmission by contact, (iii) shifting rice season later, by sowing in August or even September instead of July (the most likely mechanism being the desynchronization between rice cultivation and virus sources such as reservoir plants or vectors). Also, various farmers state their incapacity to fight the disease. More work is required to specifically address agronomic and sociological questions related to rice yellow mottle disease in Banzon.

At the within-field level, we found small scale aggregation in DAS-ELISA-positive plants, reflecting field symptom-based observations (pers. com.). This is congruent with expected major within-field secondary transmission, mechanically, through (i) direct leaf-to-leaf contact in crowded plants, (ii) contaminated hands or tools, particularly when transplanting, and applying fertilizers (these agricultural practices also enhancing direct leaf-to-leaf contact), and/or (iii) small scale vector transmission, mostly by insect’s biting and chewing mouthparts (Banwo, Alegbejo & Abo, 2004; Traore et al., 2006; Traoré et al., 2009).

Our results show that yellow mottle virus prevalence is statistically affected by agronomic factors, as well as rice cultivars. Firstly, the latter suggests potential differences in terms of resistance in the cultivars used in Banzon. Rice cultivars from Burkina Faso are moderately tolerant to various RYMV isolates experimentally. Three cultivars from Burkina Faso (FKR19, FKR62N, FKR64), commonly grown in the study sites (Barro et al., 2021a) were found to be susceptible, but less than IR64, in our experiments, which is congruent with the results obtained in the field for FKR19 and FKR64 (Sérémé et al., 2016a) and in semi-controlled experiments for FKR19 and FKR62N (Sérémé et al., 2016b). On the other hand, FKR34, commonly grown in Banzon in 2017 only, is associated with higher RYMV prevalences, which corresponds to the temporal pattern of highest disease burden in 2017. Because FKR34 was not included in our experimental infections, we cannot rule out any of the two following hypotheses: (i) FKR34 is more susceptible than other cultivars grown in Burkina Faso, or (ii) other factors led to a higher disease pressure in 2017, resulting in spurious association between FKR34 cultivar and disease prevalence. Finally, field data associating cultivar names and RYMV prevalence have to be taken with caution, as they rely on cultivar names given by the farmers, which do not always correspond to genetically well-defined cultivars, according to a recent study performed in the same study sites (Barro et al., 2021b).

Second, we tested for an effect of agronomic factors, namely off-season cultivation and the number of urea application, that were actually highly correlated, allowing to distinguish intensively cultivated fields (various urea application and two rice seasons) from low (one urea application or less, and only one rice season). We showed that this factor, in interaction with the rice cultivar, affects yellow mottle disease prevalences. This relates to the effect of fertilization on yellow mottle disease (nitrogen effect, Bouet et al., 2012), and the likely increased viral transmission associated with longer rice cultivation period (although yellow mottle disease is much less observed during the dry season; Issaka et al., 2012).

A high genetic diversity of RYMV was found in the studied site, with the co-existence of at least four genetic lineages. Over the whole period, we have three common groups (S1ca: 39.4%, S1wa: 15.2% and S1bzn: 42.4%), and a rare strain: S2 (3.0%). The genetic diversity in the studied site (2015-2019) is 0.039 sub/site. This is a very high level of diversity considering the small geographical scale of this study (the irrigated perimeter of Banzon covers ca. 520 ha), and compared to the estimates for the whole Burkina Faso (0.045 sub/site) and for West Africa (0.063 sub/site). Various West African countries have lower level of diversity than this peculiar site, and although some sampling bias (in terms of number of samples, but also in the geographic distribution of sampling within the countries) may limit this comparison, this reflects the impressive RYMV diversity at local scale in this hotspot site.

A recent study showed a relationship between the genetic composition of pathogen populations and disease intensity, with higher infection rates in wild plant populations infected by fungal pathogen populations harboring higher strain diversity (Eck et al., 2022). Following this, the high prevalence of yellow mottle disease in Banzon fields may somehow be a consequence of this very high RYMV viral diversity, but this remains speculative, especially as we have no information on the viral genetic diversity in the five other sites with lower disease pressure. The conditions that lead to yellow mottle disease outbreaks are still poorly understood (Traoré et al., 2009), and deserve further research.

Interestingly, the various lineages coexist at the rice field scale, with up to three distinct lineages detected simultaneously in delimited areas of around 625 m². Co-occurrence of various lineages at a very local scale is particularly interesting as it is a favorable context for co-infection (multiple infection of a rice plant by various RYMV genetic lineages, see below), a prerequisite for recombination.

Our results show that at least eight successful introductions of RYMV occurred in southwestern Burkina Faso including the irrigated perimeter of Banzon, enhancing the viral diversity locally. Indeed, phylogeographic reconstructions suggest that strains S1wa and S2 were introduced first, most likely during the 1980s, while the strain S1ca arrived later, probably after 2000. The irrigated perimeter of Banzon was built in early 1980s’ (Toe, 1992), as well as others in the region (Nebie, 1993). In the last decades, Burkina Faso experienced rice cultivation intensification (yield doubled between 1960’s and 1980’s, FAO), followed by an increase in areas cultivated in rice (fourfold increase between the period before and after 2018, FAO). Increased exchanges of material, and possibly highest probability of new introduction, are likely associated with these recent changes in rice cultivation. Within one irrigated site, we found the coexistence of genetic groups introduced decades ago and others recently emerged. Recombination event between S1wa and S1ca involving the CP coding region, either occurring locally (in Banzon site) or elsewhere (in the area), led to the recent emergence of a lineage (S1bzn), maintained over the 5-year period. Other studies on crop viruses have documented the emergence of recombinant groups with variable outcomes. For example, a recombinant strain of pepino mosaic virus (PepMV) found in 2004 may be extinct as it was not detected again since 2005 (Alcaide et al., 2020), while for tomato yellow leaf curl virus (TYLC), a recombinant strain displaced its parental viruses in southern Morocco (Belabess et al., 2015). The success of the recombinant strain of TYLC in Morocco is likely the consequence of a selective advantage, as it exhibits greater within-host viral accumulation in a resistant cultivar that had been recently deployed at the time of emergence (Belabess et al., 2016).

The source of these introductions may be the neighboring rainfed lowlands. However, we notice that the studied site in the neighborhood (Banzon-RL= Senzon village) experiments low level of virus circulation. Alternatively, long-distance migration events may explain observed patterns, in accordance with the average dispersal rate of 13 km/year estimated for RYMV in West Africa (Trovão et al., 2015). Putative transmission events may involve insect vectors flying long distances, or the transport of infective tissue (leaf fragments or empty rice spikelets) associated with farmers seed exchanges (Banwo, Alegbejo & Abo, 2004). Significant level of exchange of infected material between farmers from distant areas in Burkina Faso is expected considering that some rice pathogens exhibit low level of spatial structure in this country (Kaboré et al., 2022). Control measures would include a more careful control of rice material imports to prevent introduction of new pathogen strains. The high genetic diversity of RYMV in Banzon illustrates the increased viral circulation from the 1980s that gradually blurs the strong geographical structure of RYMV documented at the continental level (Traore et al., 2005; Dellicour et al., 2018).

Although little geographic structure was found within the studied irrigated perimeter, we detected genetic differentiation between the 5 sampling years, and temporal variation in frequencies of the genetic groups. A recently introduced group, S1bzn, seems to increase in frequency over the studied period, from 14% in 2015 to 68% in 2019, while the strain S1ca, particularly frequent at the beginning of the survey (76% in 2015) was found at only 14% in 2018 and 32% in 2019. The S1wa lineage was not found in 2019, but we hypothesize it is not extinct as it was common in the previous year (64% in 2018). The rare S2 strain was not found in the last two surveyed years, either because it was not sampled by chance, or because it was actually absent these years and maybe now extinct.

The turnover rate of RYMV genetic groups over the study period raises questions on the underlying processes, including annual founding effects or selective pressures. To investigate the selection hypothesis, we characterized as a fitness proxy the outcome of infection by viral isolates belonging to the three genetic groups (S1wa, S1ca and S1bzn) under controlled conditions. The experimental results showed that induced symptoms and viral load varied between isolates, while the genetic group was not a determinant of infection outcome. These results are congruent with other studies in plant viruses, where the genetic groups are not homogeneous in terms of pathogenicity (see Fraile et al., 2011; N’Guessan et al., 2000).

However, interestingly, we found that the isolate with the highest viral load and 3 out of the 5 most aggressive isolates belong to the S1bzn genetic lineage, which is in agreement with the hypothesis of a selective advantage of the lineage S1bzn (Figure 6). Such an effect of natural selection is supported by the overall increase of this lineage through years (Figure 5). However, our experimental fitness estimates may have been too simple and considering the outcome of co-infections by various RYMV strains would be interesting in further experiments. Indeed, in Ivory Coast, although no difference between two strains could be detected in mono-infections, the S2 strain of RYMV from Ivory Coast dominates the S1 strain in experimental co-infection (N’Guessan et al., 2000). Multiple RYMV infections may be quite common in western Burkina Faso, as we obtained mixed Sanger chromatograms for 4.3% of sequenced samples. Long-term coexistence of two viral strains may occur in spite of differential fitness in single infection, provide that the same strain is not favored both in mono-infections and in mixed infections, as exemplified for TYLC in La Reunion (Péréfarres et al., 2014).

Innovations in science and technology, including outbreak forecasting, constitute a promise for improving the management of viral epidemics, but this requires detailed understanding of small-scale epidemiology. For the rice yellow mottle virus, Hébrard et al., (2018) built a resistance-breaking risk map at the scale of the African continent, based on the geographical repartition of the capacity of isolates to evolve and infect cultivars with resistance genes and alleles in controlled conditions. This work also identified a hypervirulent pathotype within the S1ca group, able to overcome all the sources of high resistance identified in rice. Our work focusing on the local scale (region, site and field level) in western Burkina Faso illustrates the complexity of durable resistance deployment, as multiple genetic groups, likely presenting different resistance-breaking abilities, co-exist in the same site. This highlights the importance to track the different RYMV pathotypes with innovative and effective detection tools. Further, it urges to continue the development of new resistances, including 1) the combination of multiple resistance genes into a single plant genotype (pyramiding), and 2) the introduction of innovative way to create new resistance (such as CRISPR). This monitoring will be followed up as they evidence the relevance of this pathosystem to address major current questions for the ecology and evolution of plant viruses, including the impact of multiple infections and recombination in viral evolution (Lefeuvre et al., 2019).

## ACKNOWLEDGEMENTS

We thank Laurence Albar and Agnès Pinel-Galzi for constructive discussions, as well as Emilie Thomas, Manon Perez and Pascaline Boyer for help in the experimental work. We are grateful to Sylvain Zougrana, Kader Guigma, Edouard Kabore, Yacouba Kone, Martial Kabore, Jean-Noël Ouattara, Moumouni Traoré and Roméo Dabiré contributed to the sampling in southwestern Burkina Faso. We thank rice farmers from all the study sites (Badala, Bama, Banzon, Karfiguela, Senzon and Tengrela) for their collaboration.

This work was publicly funded through ANR (the French National Research Agency) under the program “Investissements d’avenir” with the reference ANR-10-LABX-001-01 Labex Agro (VIRBIARF, E- Space and RiPaBIOME projects), coordinated by Agropolis Fondation under the frame of I-SITE MUSE (ANR-16-IDEX-006) and by the CGIAR Research Program on Rice Agri-food Systems (RICE).

