## Supplementary material for "Dynamics of the rice yellow mottle disease in western Burkina Faso: epidemic monitoring, spatio-temporal variation of viral diversity and pathogenicity in a disease hotspot"

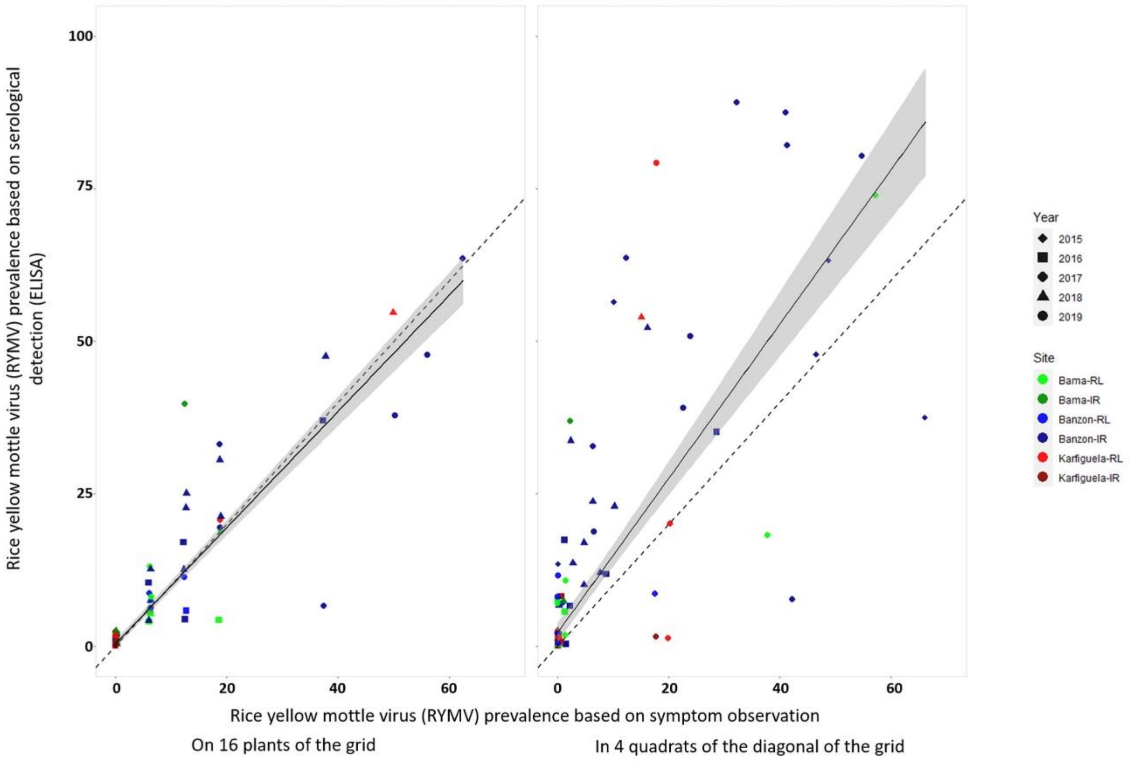


**Supplementary Figure 1:** Relationship between RYMV prevalence based on symptom observations and RYMV prevalence based on serological diagnostic results.

Each point corresponds to a particular observation (one field and year). Dotted line corresponds to the bisector (slope = 1). Solid line was obtained using the “stat_smooth” function in R.
The two graphs have the same variable (prevalence based on RYMV-specific DAS-ELISA results) on the y-axis. On the other hand, x-axis represent for both graphs the prevalence based on yellow mottle disease symptom observations but based on the observations of the 16 plants of the grid on the left (same plants analyzed using DAS-ELISA) while on the right are represented the prevalence based on visual inspection of the four diagonal quadrats.

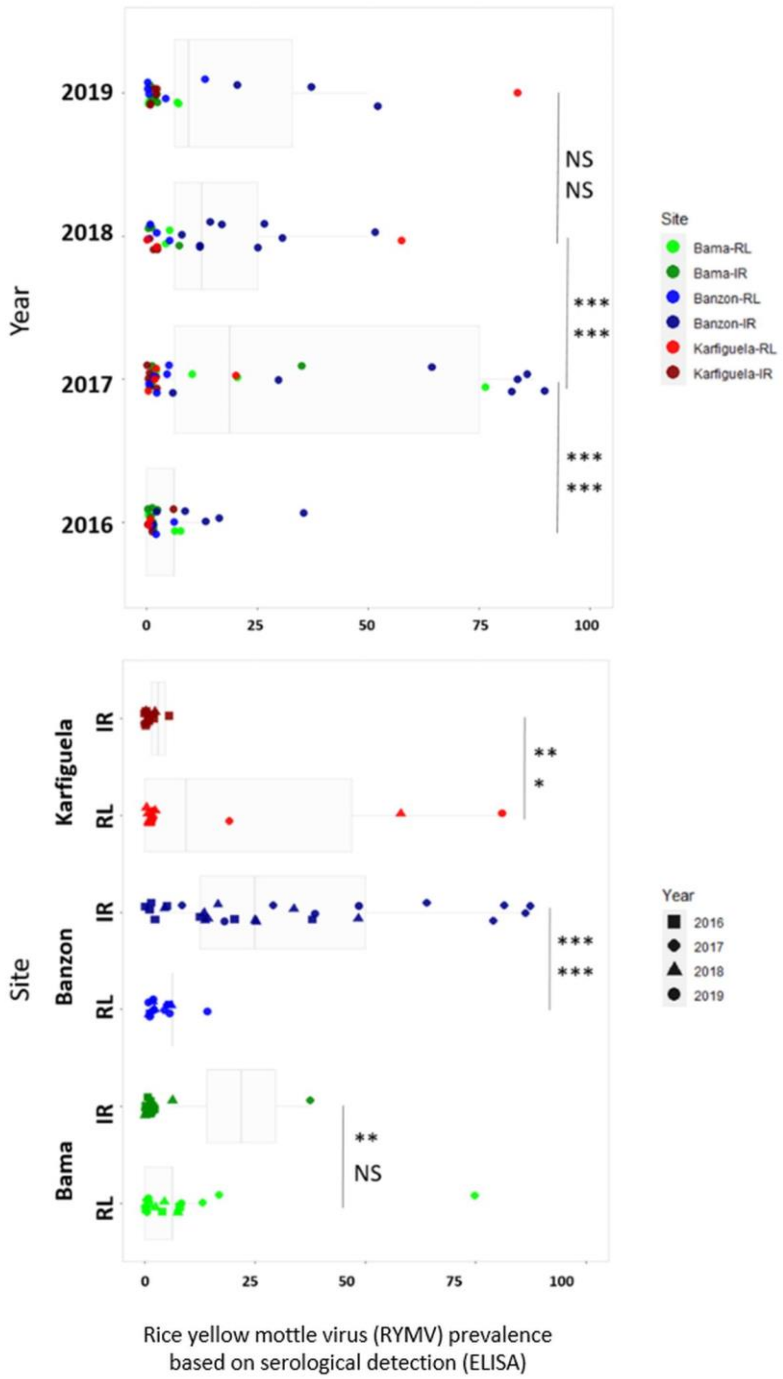


**Supplementary Figure 2:** Temporal and spatial variation in yellow mottle disease prevalence over all the study sites in the 2016-2019 period.

Each point is one observation (179 over the 2016-2019 period). Only the case where yellow mottle disease was observed (prevalence higher than zero) were used for boxplots. Statistical results are reported both including (on top) or not (on bottom) the cases where no yellow mottle disease could be found.

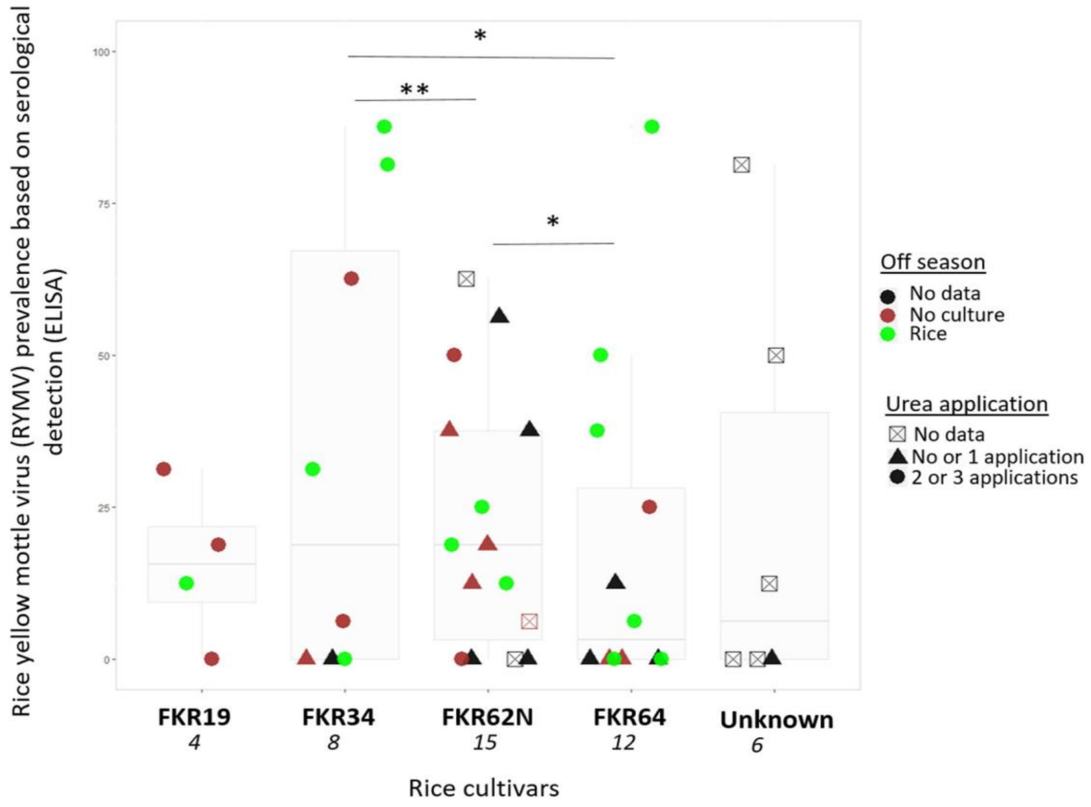


**Supplementary Figure 3:** Relationship between RYMV prevalence (based on serological detection) and agricultural practices (named: off season land use, urea application) within the irrigated perimeter of Banzon. Asterisks represent significant differences between rice cultivars (Tukey honestly significant difference, * $p < 0.05$ and ** $p < 0.001$).

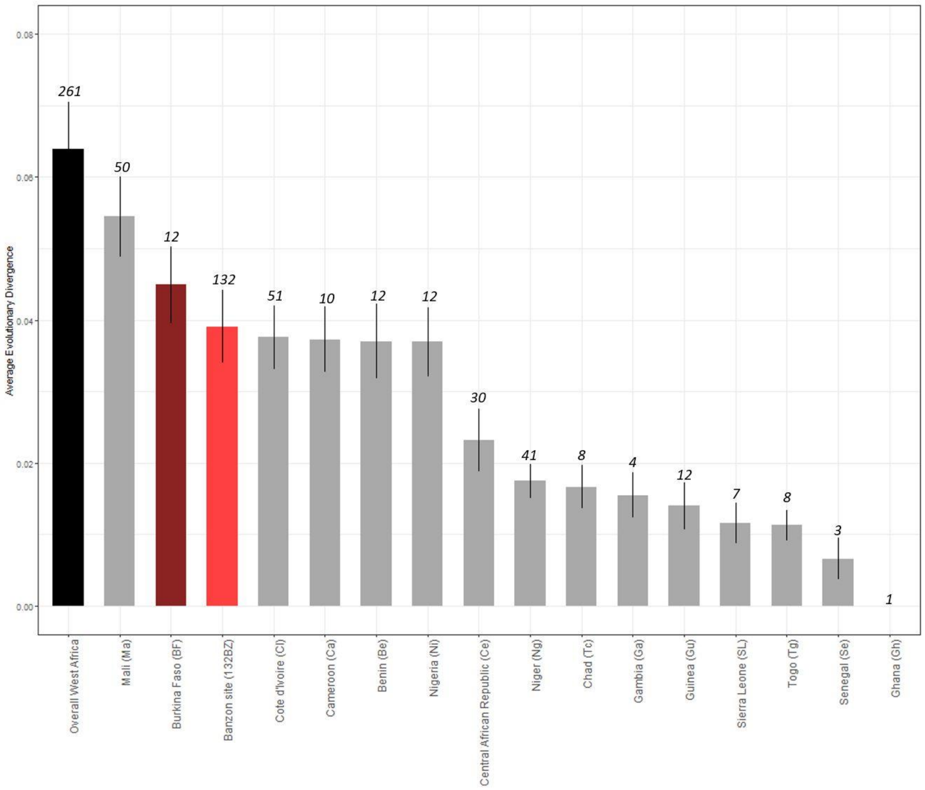


**Supplementary Figure 4**: Genetic diversity (d) of RYMV populations circulating in West- and West-Central Africa (Overall, 261CP) and specifically in Benin (Be), Burkina Faso (BF), Cameroon (Ca), Central African Republic (Ce), Chad (Ch), Ivory Coast (CI), Ghana (Gh), Guinea (Gu), Mali (Ma), Niger (Ng), Nigeria (Ni), Togo (Tg) and in Banzon locality (132BZ). S.E.: standard errors; nd: not determined.

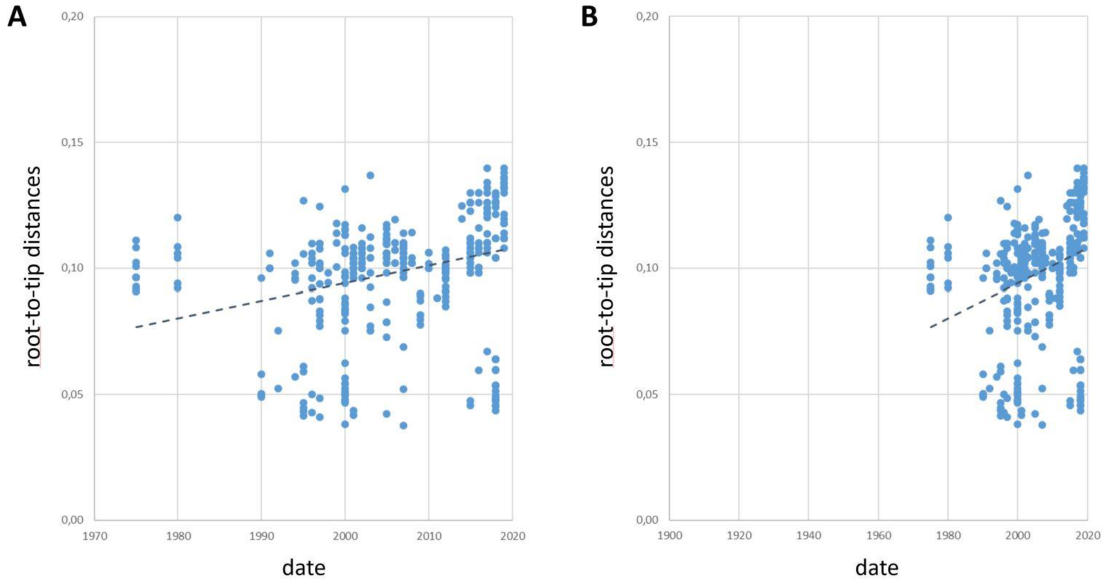


**Supplementary Figure 5**: Exploration of the temporal signal from the RYMV 393 CP dataset by linear regression approach. Root-to-tip divergence as a function of sampling time for Maximum Likelihood tree clusters obtained with 393 non-recombinant RYMV CP sequences. Plot representations in a 1970-2020 (A) and a 1900-2020 (B) dating range.

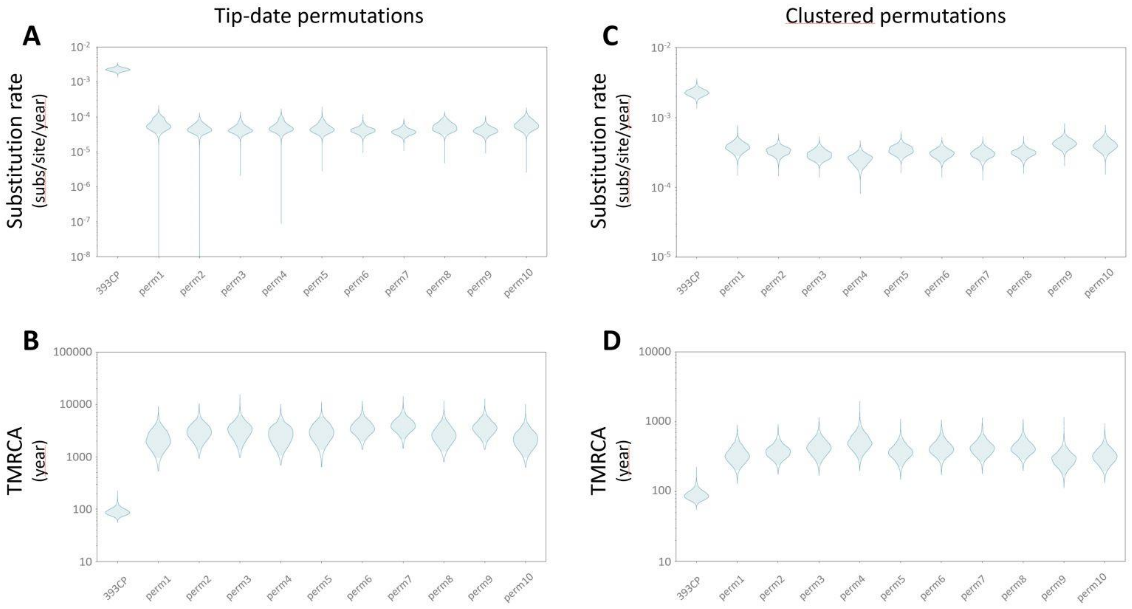


**Supplementary Figure 6**: Exploration of the temporal signal from the RYMV 393 CP dataset by date-randomization tests. Violin plots representing the intervals of the substitution rate (substitution/site/year; A and C) and the time to most recent common ancestor (years; B and D) in the RYMV CP sequence dataset from the sampled dates (393 CP) and from the dates after tip (A and B) or cluster (C and D) permutations (from perm1 to perm10). The blue areas represent the range of the 95% HPD estimates.

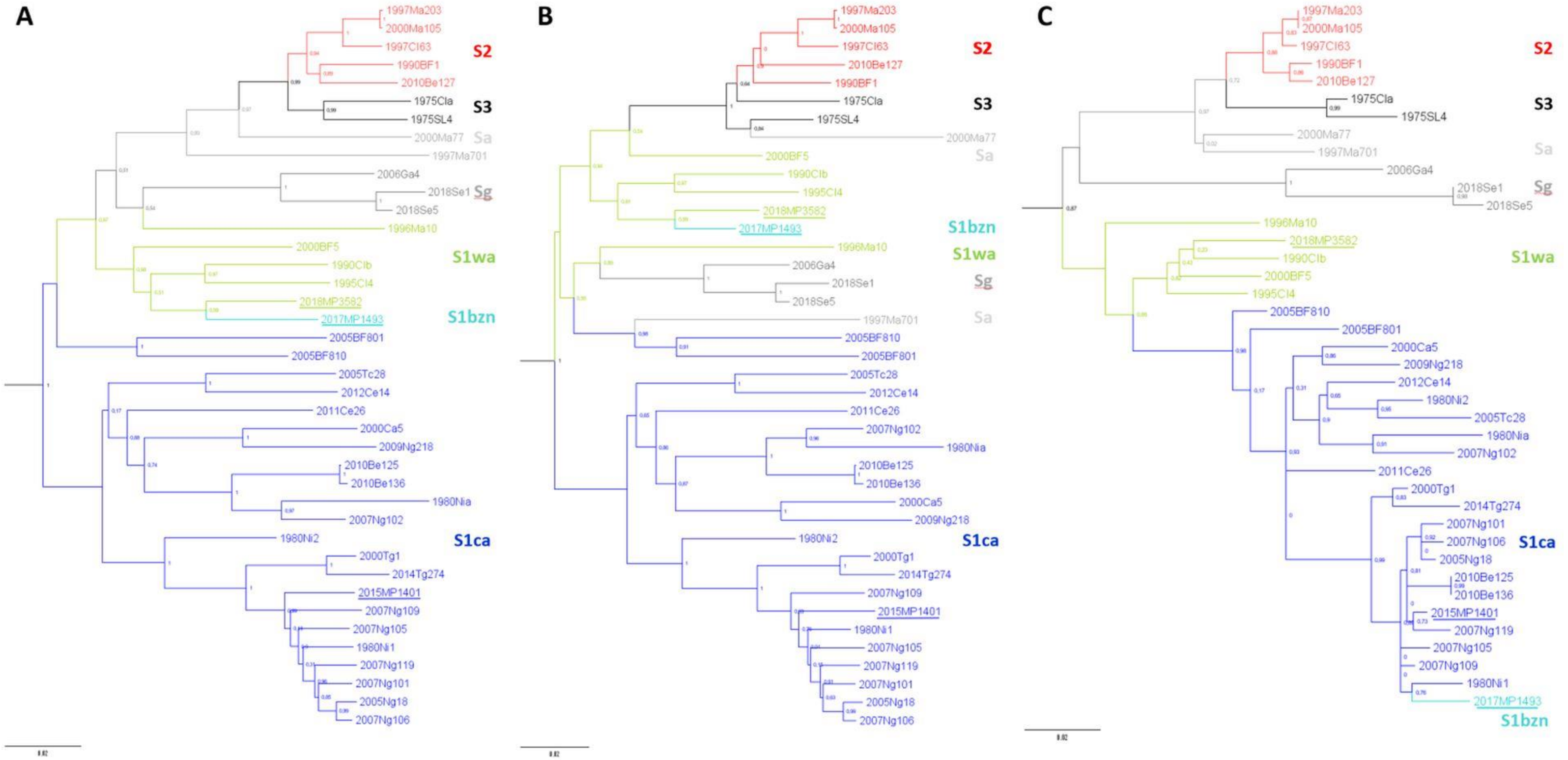


Supplementary Figure 7: Maximum-Likelihood (ML) phylogenetic trees of 37 RYMV reference isolates from West- and West-Central Africa (from the databases) and three isolates from Banzon (this study).

The ML phylogenetic trees were built based on the full-length genome sequences, 4461 nt (A), RYMV full-length genomes without the recombinant domains corresponding to the coat protein (CP), 3743 nt (B) and only CP sequences, 722 nt (C). The different colors correspond to the identified genetic groups: S2 (red), S3 (dark), Sa (light grey), Sg (dark grey), S1wa (green), S1ca (dark blue) and S1bzn (light blue). The three isolates from Banzon (2015MP1401, 2017MP1493, 2018MP3582), are underlined.

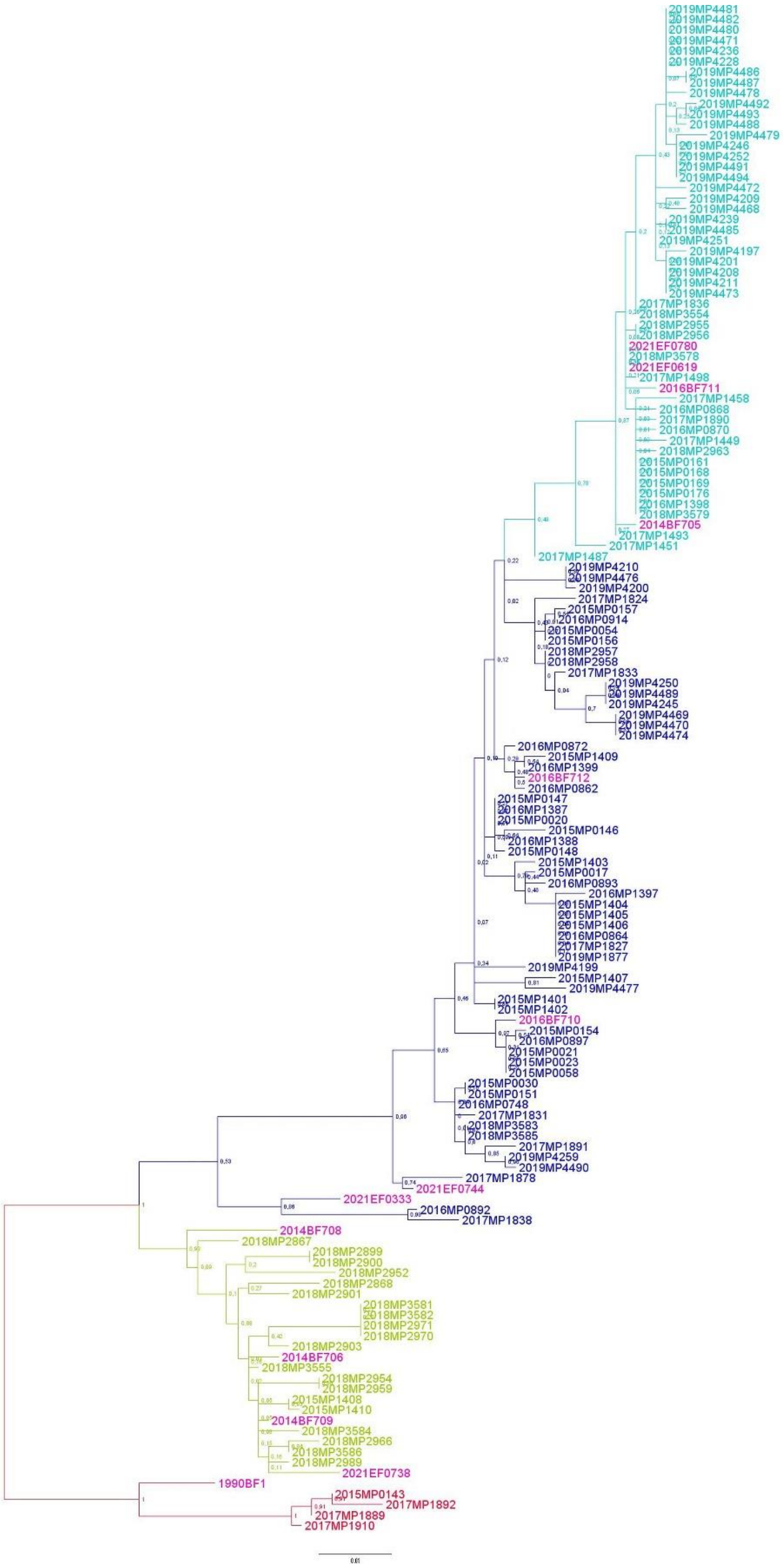


**Supplementary Figure 8**: Phylogenetic tree of coat protein (CP) sequences of RYMV samples collected between 2015 and 2019 within the irrigated area of Banzon and 13 isolates used in the experimental part (in pink). The colors represent the four identified genetic groups: S2 in red, S1wa in green, S1ca in dark blue and the recombinant line S1bzn in light blue.

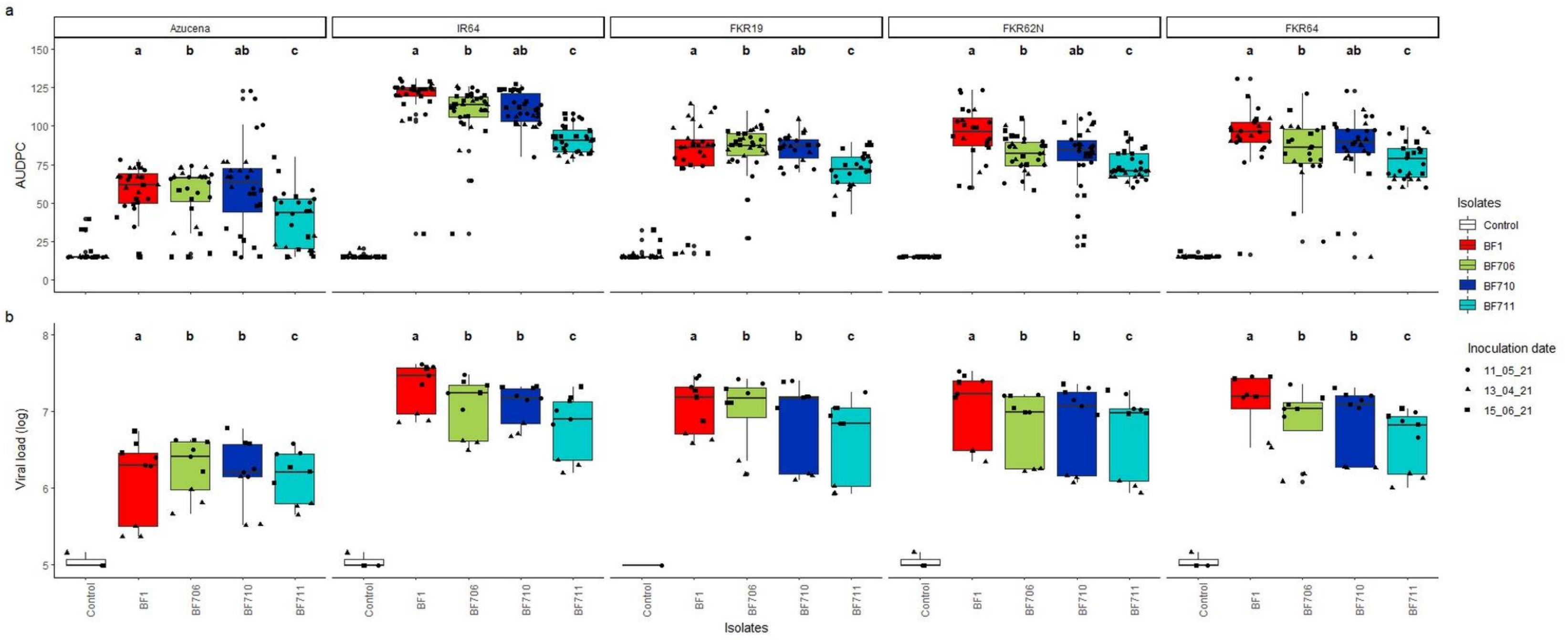


**Supplementary Figure 9**: Symptom estimate and viral load of four RYMV isolates from the four genetic groups identified in Banzon (S2, S1wa, S1ca, S1bzn) infecting five distinct rice cultivars (three commonly grown in Burkina Faso and two reference cultivars).

a. The symptom severity estimated by the area under disease progress curve (AUDPC) of the four RYMV isolates inoculated on the five rice cultivars. AUDPC was determined according the notation severity scale over five dates (from 9 to 24 dpi), on 45 phenotyped plants.

b. The intra-plant viral load of the four RYMV isolates for each of the five rice cultivars. The number of RYMV copies was estimated by RT-qPCR on 9 meristematic area collected at 7 dpi using a standard curve.

RYMV isolates were inoculated on three rice cultivars from Burkina Faso (FKR19, FKR62N, FKR64) and partially resistant (Azucena) and susceptible (IR64) reference rice cultivars. The four RYMV isolates are represented by four different colors corresponding to their genetic group, S2 (red), S1wa (green), S1ca (dark blue) and S1bzn (light blue). Three repetitions were performed represented by three dot shapes (inoculation dates 13/04/21; 11/05/21; 15/06/21) and letters indicate the significant differences between each isolate (Tukey honestly significant difference $p < 0.05$).

**Supplementary Table 1**: Statistical analyses of the RYMV prevalence based on serological detection.

Results of the GLM including 'Year' and 'Rice system*Zone' are reported. On the left, are shown the results on GLM on the whole dataset, including the fields where no yellow mottle symptoms were observed while on the right are only the cases where yellow mottle symptoms were observed. Contrast analysis were performed for the effect of 'Year', and for the effect of 'Rice system' within each geographical zone.

| | All dataset (179 observations) | | | Only fields with RYMV (54 observations) | | |
|---|---|---|---|---|---|---|
| Model (GLM) | | | | | | |
| | Chisq | Df | Pr(>Chisq) | Chisq | Df | Pr(>Chisq) |
| Year | 85.406 | 3 | <.0001 | 71.984 | 3 | <.0001 |
| Rice_system | 11.271 | 1 | 0.0008 | 1.351 | 1 | 0.2451 |
| Zone | 271.432 | 2 | <.0001 | 23.708 | 2 | <.0001 |
| Rice_system:Zone | 154.310 | 2 | <.0001 | 34.256 | 2 | <.0001 |
| Temporal variation | | | | | | |
| | estimate | z.ratio | p.value | estimate | z.ratio | p.value |
| 2016-2017 | -2.144 | -7.355 | <.0001 | -2.108 | -6.964 | <.0001 |
| 2016-2018 | -0.808 | -2.629 | 0.0426 | -0.829 | -2.612 | 0.0445 |
| 2016-2019 | -1.105 | -3.444 | 0.0032 | -1.454 | -4.282 | 0.0001 |
| 2017-2018 | 1.336 | 6.372 | <.0001 | 1.279 | 5.627 | <.0001 |
| 2017-2019 | 1.040 | 4.636 | <.0001 | 0.655 | 2.648 | 0.0403 |
| 2018-2019 | -0.296 | -1.191 | 0.6328 | -0.624 | -2.256 | 0.1085 |
| Rice system effect | | | | | | |
| Contrats IR-RL | estimate | z.ratio | p.value | estimate | z.ratio | p.value |
| BAMA | -1.35 | 3.081 | 0.0021 | 0.601 | -1.197 | 0.2314 |
| BANZON | 3.18 | -7.944 | <.0001 | 2.172 | -5.180 | <.0001 |
| KARFIGUELA | -3.53 | 3.446 | 0.0006 | -2.450 | 2.316 | 0.0206 |

**Supplementary Table 2**: Contingency table of the frequency distribution of the four CP genetic groups over time, from each field (5 sampled fields within the irrigated perimeter of Banzon) and overall (the five fields considered together).

| | **2015** | | | | | | **2016** | | | | | | **2017** | | | | | | **2018** | | | | | | **2019** | | | | | |
|---|---|---|---|---|---|---|---|---|---|---|---|---|---|---|---|---|---|---|---|---|---|---|---|---|---|---|---|---|---|---|
| | BZ02 | BZ04 | BZ09 | BZ10 | BZ11 | **all** | BZ02 | BZ04 | BZ09 | BZ10 | BZ11 | **all** | BZ02 | BZ04 | BZ09 | BZ10 | BZ11 | **all** | BZ02 | BZ04 | BZ09 | BZ10 | BZ11 | **all** | BZ02 | BZ04 | BZ09 | BZ10 | BZ11 | **all** |
| **S1ca** | 5 | 4 | 5 | 8 | 0 | **22** | 3 | 0 | 3 | 3 | 3 | **12** | 0 | 5 | 2 | 1 | 0 | **8** | 0 | 0 | 2 | 2 | 0 | **4** | 0 | 7 | 0 | 1 | 5 | **13** |
| **S1bzn** | 0 | 0 | 0 | 0 | 4 | **4** | 0 | 0 | 2 | 0 | 1 | **3** | 0 | 1 | 3 | 1 | 3 | **8** | 1 | 0 | 3 | 2 | 0 | **6** | 0 | 9 | 0 | 9 | 10 | **28** |
| **S1wa** | 0 | 0 | 0 | 2 | 0 | **2** | 0 | 0 | 0 | 0 | 0 | **0** | 0 | 0 | 0 | 0 | 0 | **0** | 3 | 4 | 4 | 6 | 1 | **18** | 0 | 0 | 0 | 0 | 0 | **0** |
| **S2** | 0 | 0 | 1 | 0 | 0 | **1** | 0 | 0 | 0 | 0 | 0 | **0** | 0 | 0 | 0 | 2 | 1 | **3** | 0 | 0 | 0 | 0 | 0 | **0** | 0 | 0 | 0 | 0 | 0 | **0** |

**Supplementary Table 3**: Information on the 13 RYMV isolates used in the two experimental infections (isolates used for phenotyping).

| ISOLATE | SAMPLING YEAR | SITE | RICE SYSTEM | LATITUDE | LONGITUDE | GENETIC LINEAGE | REFERENCE Genebank Accession number |
|---|---|---|---|---|---|---|---|
| **BF1** | 1997 | | | | | S2 | Pinel et al., 2000<br>Accession no : AJ279901 |
| **BF705** | 2014 | Banzon | Irrigated | 11.31955 | -4.80978 | S1bzn | Tollenaere et al., 2017<br>Accession no : (to be assigned) |
| **BF706** | 2014 | Karankasso Sambla | Rainfed lowland | 11.24732 | -4.56256 | S1wa | Tollenaere et al., 2017<br>Accession no : (to be assigned) |
| **BF708** | 2014 | Banfora | Rainfed lowland | 10.63019 | -4.77723 | S1wa | This study<br>Accession no : (to be assigned) |
| **BF709** | 2014 | Banzon | Irrigated | 11.31955 | -4.80978 | S1wa | This study<br>Accession no : (to be assigned) |
| **BF710** | 2016 | Banzon | Irrigated | 11.321635 | -4.805467 | S1ca | This study<br>Accession no : (to be assigned) |
| **BF711** | 2016 | Banzon | Irrigated | 11.325212 | -4.807832 | S1bzn | This study<br>Accession no : (to be assigned) |
| **BF712** | 2016 | Banzon | Irrigated | 11.329952 | -4.801317 | S1ca | This study<br>Accession no : (to be assigned) |
| **EF0333** | 2021 | Badala | Rainfed lowland | 11.387242 | -4.363315 | S1ca | This study<br>Accession no : (to be assigned) |
| **EF0619** | 2021 | Banzon | Irrigated | 11.33554 | -4.80248 | S1bzn | This study<br>Accession no : (to be assigned) |
| **EF0738** | 2021 | Banzon | Irrigated | 11.32138 | -4.803342 | S1wa | This study<br>Accession no : (to be assigned) |
| **EF0744** | 2021 | Banzon | Irrigated | 11.323422 | -4.803110 | S1ca | This study<br>Accession no : (to be assigned) |
| **EF0780** | 2021 | Banzon | Irrigated | 11.323422 | -4.803110 | S1bzn | This study<br>Accession no : (to be assigned) |

**Supplementary Table 4**: Contrast analysis for Experiment 1 and Experiment 2 of the phenotypic evaluation of RYMV isolates: estimations of symptom severity (AUDPC) and viral load.

Contrast analyses were performed for the effect of 'Rice cultivars' in the Experiment 1 and for the effect of 'Isolates' in both experiments. Tukey HSD results on symptom severity (AUDPC data) are shown on the left and on viral load on the right, for each experiment.

**Experiment 1**

| Tukey HSD | AUDPC | | | Viral load | | |
|---|---|---|---|---|---|---|
| Rice cultivar | estimate | t.ratio | p.value | estimate | t.ratio | p.value |
| Azucena - IR64 | -53.98 | -23.701 | <.0001 | -13550407 | -11.416 | <.0001 |
| Azucena - FKR19 | -27.03 | -11.038 | <.0001 | -9787714 | -8.181 | <.0001 |
| Azucena - FKR62N | -29.61 | -12.411 | <.0001 | -9357547 | -7.883 | <.0001 |
| Azucena - FKR64 | -31.99 | -12.971 | <.0001 | -8353554 | -6.927 | <.0001 |
| IR64 - FKR19 | 26.94 | 11.255 | <.0001 | 3762693 | 3.145 | 0.017 |
| IR64 - FKR62N | 24.36 | 10.483 | <.0001 | 4192860 | 3.532 | 0.005 |
| IR64 - FKR64 | 21.98 | 9.112 | <.0001 | 5196853 | 4.310 | 0.0003 |
| FKR19 - FKR62N | -2.58 | -1.086 | 0.814 | 430167 | 0.360 | 0.996 |
| FKR19 - FKR64 | -4.96 | -2.028 | 0.254 | 1434161 | 1.181 | 0.762 |
| FKR62N - FKR64 | -2.38 | -0.996 | 0.857 | 1003993 | 0.833 | 0.920 |

| Isolate | estimate | t.ratio | p.value | estimate | t.ratio | p.value |
|---|---|---|---|---|---|---|
| BF1 - BF706 | 5.72 | 2.702 | 0.036 | 5147974 | 4.760 | <.0001 |
| BF1 - BF710 | 4.03 | 1.879 | 0.238 | 5012911 | 4.692 | <.0001 |
| BF1 - BF711 | 17.97 | 8.443 | <.0001 | 8537045 | 7.991 | <.0001 |
| BF706 - BF710 | -1.69 | -0.814 | 0.848 | -135063 | -0.126 | 0.9993 |
| BF706 - BF711 | 12.25 | 5.926 | <.0001 | 3389072 | 3.152 | 0.010 |
| BF710 - BF711 | 13.95 | 6.655 | <.0001 | 3524135 | 3.319 | 0.006 |

**Experiment 2**

| Tukey HSD | AUDPC | | | Viral load | | |
|---|---|---|---|---|---|---|
| Isolate | estimate | t.ratio | p.value | estimate | t.ratio | p.value |
| BF1 S2 - EF0619 S1bzn | -9.8029 | -4.165 | 0.003 | -12338620 | -5.291 | 0.0001 |
| BF1 S2 - EF0780 S1bzn | -0.0342 | -0.015 | 1.000 | 2126880 | 0.912 | 0.999 |
| BF1 S2 - BF705 S1bzn | 1.700 | 0.734 | 0.999 | 2409627 | 1.033 | 0.998 |
| BF1 S2 - BF711 S1bzn | 40.700 | 17.573 | <.0001 | 3215517 | 1.379 | 0.975 |
| BF1 S2 - BF710 S1ca | 6.151 | 2.567 | 0.335 | 3049888 | 1.308 | 0.984 |
| BF1 S2 - EF0744 S1ca | 14.017 | 5.904 | <.0001 | 3109815 | 1.333 | 0.981 |
| BF1 S2 - BF712 S1ca | 27.128 | 11.200 | <.0001 | 3195256 | 1.370 | 0.976 |
| BF1 S2 - EF0333 S1ca | 31.924 | 13.560 | <.0001 | 3196809 | 1.371 | 0.976 |
| BF1 S2 - BF706 S1wa | 13.714 | 5.826 | <.0001 | 2783323 | 1.193 | 0.992 |
| BF1 S2 - EF0738 S1wa | 16.897 | 7.238 | <.0001 | 3195302 | 1.370 | 0.976 |
| BF1 S2 - BF708 S1wa | 21.815 | 9.184 | <.0001 | 3211058 | 1.377 | 0.975 |
| BF1 S2 - BF709 S1wa | 56.888 | 24.563 | <.0001 | 3208614 | 1.376 | 0.975 |
| EF0619 S1bzn - EF0780 S1bzn | 9.769 | 4.183 | 0.003 | 14465501 | 6.203 | <.0001 |
| EF0619 S1bzn - BF705 S1bzn | 11.503 | 4.925 | 0.0001 | 14748248 | 6.324 | <.0001 |

| | | | | | | |
|---|---|---|---|---|---|---|
| EF0619 S1bzn - BF711 S1bzn | 50.503 | 21.624 | <.0001 | 15554138 | 6.669 | <.0001 |
| EF0619 S1bzn - BF710 S1ca | 15.954 | 6.606 | <.0001 | -15388509 | -6.598 | <.0001 |
| EF0619 S1bzn - EF0744 S1ca | 23.820 | 9.953 | <.0001 | -15448435 | -6.624 | <.0001 |
| EF0619 S1bzn - BF712 S1ca | 36.931 | 15.136 | <.0001 | -15533876 | -6.661 | <.0001 |
| EF0619 S1bzn - EF0333 S1ca | 41.727 | 17.583 | <.0001 | -15535429 | -6.661 | <.0001 |
| EF0619 S1bzn - BF706 S1wa | 23.517 | 9.911 | <.0001 | -15121943 | -6.484 | <.0001 |
| EF0619 S1bzn - EF0738 S1wa | 26.700 | 11.344 | <.0001 | -15533922 | -6.661 | <.0001 |
| EF0619 S1bzn - BF708 S1wa | 31.618 | 13.208 | <.0001 | -15549678 | -6.668 | <.0001 |
| EF0619 S1bzn - BF709 S1wa | 66.691 | 28.555 | <.0001 | -15547234 | -6.667 | <.0001 |
| EF0780 S1bzn - BF705 S1bzn | 1.734 | 0.755 | 0.9999 | 282747 | 0.121 | 1.000 |
| EF0780 S1bzn - BF711 S1bzn | 40.734 | 17.730 | <.0001 | 1088637 | 0.467 | 1.000 |
| EF0780 S1bzn - BF710 S1ca | 6.185 | 2.601 | 0.3136 | -923008 | -0.396 | 1.000 |
| EF0780 S1bzn - EF0744 S1ca | 14.052 | 5.964 | <.0001 | -982935 | -0.421 | 1.000 |
| EF0780 S1bzn - BF712 S1ca | 27.163 | 11.300 | <.0001 | -1068376 | -0.458 | 1.000 |
| EF0780 S1bzn - EF0333 S1ca | 31.959 | 13.682 | <.0001 | -1069928 | -0.459 | 1.000 |
| EF0780 S1bzn - BF706 S1wa | 13.748 | 5.887 | <.0001 | -656443 | -0.281 | 1.000 |
| EF0780 S1bzn - EF0738 S1wa | 16.932 | 7.311 | <.0001 | -1068421 | -0.458 | 1.000 |
| EF0780 S1bzn - BF708 S1wa | 21.849 | 9.270 | <.0001 | -1084177 | -0.465 | 1.000 |
| EF0780 S1bzn - BF709 S1wa | 56.922 | 24.775 | <.0001 | -1081734 | -0.464 | 1.000 |
| BF705 S1bzn - BF711 S1bzn | 39.000 | 16.975 | <.0001 | 805890 | 0.346 | 1.000 |
| BF705 S1bzn - BF710 S1ca | 4.451 | 1.871 | 0.8110 | -640261 | -0.275 | 1.000 |
| BF705 S1bzn - EF0744 S1ca | 12.317 | 5.227 | <.0001 | -700188 | -0.300 | 1.000 |
| BF705 S1bzn - BF712 S1ca | 25.428 | 10.579 | <.0001 | -785629 | -0.337 | 1.000 |
| BF705 S1bzn - EF0333 S1ca | 30.224 | 12.940 | <.0001 | -787181 | -0.338 | 1.000 |
| BF705 S1bzn - BF706 S1wa | 12.014 | 5.144 | <.0001 | -373696 | -0.160 | 1.000 |
| BF705 S1bzn - EF0738 S1wa | 15.197 | 6.562 | <.0001 | -785674 | -0.337 | 1.000 |
| BF705 S1bzn - BF708 S1wa | 20.115 | 8.535 | <.0001 | -801430 | -0.344 | 1.000 |
| BF705 S1bzn - BF709 S1wa | 55.188 | 24.021 | <.0001 | -798987 | -0.343 | 1.000 |
| BF711 S1bzn - BF710 S1ca | -34.549 | -14.526 | <.0001 | 6903 | 0.003 | 1.000 |
| BF711 S1bzn - EF0744 S1ca | -26.683 | -11.324 | <.0001 | 105702 | 0.045 | 1.000 |
| BF711 S1bzn - BF712 S1ca | -13.572 | -5.646 | <.0001 | 20261 | 0.009 | 1.000 |
| BF711 S1bzn - EF0333 S1ca | -8.776 | -3.757 | 0.0124 | 18709 | 0.008 | 1.000 |
| BF711 S1bzn - BF706 S1wa | -26.987 | -11.555 | <.0001 | 432194 | 0.185 | 1.000 |
| BF711 S1bzn - EF0738 S1wa | -23.803 | -10.277 | <.0001 | 20216 | 0.009 | 1.000 |
| BF711 S1bzn - BF708 S1wa | -18.885 | -8.013 | <.0001 | 4460 | 0.002 | 1.000 |
| BF711 S1bzn - BF709 S1wa | 16.1875 | 7.046 | <.0001 | 6903 | 0.003 | 1.000 |
| BF710 S1ca - EF0744 S1ca | 7.8662 | 3.230 | 0.0681 | 59927 | 0.026 | 1.000 |
| BF710 S1ca - BF712 S1ca | 20.9770 | 8.460 | <.0001 | 145368 | 0.062 | 1.000 |
| BF710 S1ca - EF0333 S1ca | 25.7731 | 10.673 | <.0001 | 146920 | 0.063 | 1.000 |
| BF710 S1ca - BF706 S1wa | 7.563 | 3.131 | 0.0900 | 266565 | 0.114 | 1.000 |
| BF710 S1ca - EF0738 S1wa | 10.746 | 4.485 | 0.0007 | -145413 | -0.062 | 1.000 |
| BF710 S1ca - BF708 S1wa | 15.664 | 6.433 | <.0001 | -161169 | -0.069 | 1.000 |
| BF710 S1ca - BF709 S1wa | 50.736 | 21.331 | <.0001 | -158726 | -0.068 | 1.000 |
| EF0744 S1ca - BF712 S1ca | 13.111 | 5.328 | <.0001 | 85441 | 0.037 | 1.000 |
| EF0744 S1ca - EF0333 S1ca | 17.907 | 7.480 | <.0001 | 86993 | 0.037 | 1.000 |
| EF0744 S1ca - BF706 S1wa | -0.304 | -0.127 | 1.0000 | 326492 | 0.140 | 1.000 |
| EF0744 S1ca - EF0738 S1wa | 2.880 | 1.213 | 0.9922 | -85487 | -0.037 | 1.000 |
| EF0744 S1ca - BF708 S1wa | 7.798 | 3.229 | 0.0682 | -101242 | -0.043 | 1.000 |

| | | | | | | |
|---|---|---|---|---|---|---|
| EF0744 S1ca - BF709 S1wa | 42.870 | 18.194 | <.0001 | -98799 | -0.042 | 1.000 |
| BF712 S1ca - EF0333 S1ca | 4.796 | 1.967 | 0.7540 | 1552 | 0.001 | 1.000 |
| BF712 S1ca - BF706 S1wa | -13.415 | -5.498 | <.0001 | 411933 | 0.177 | 1.000 |
| BF712 S1ca - EF0738 S1wa | -10.231 | -4.227 | 0.0021 | -46 | 0.000 | 1.000 |
| BF712 S1ca - BF708 S1wa | -5.313 | -2.161 | 0.6200 | -15802 | -0.007 | 1.000 |
| BF712 S1ca - BF709 S1wa | 29.759 | 12.381 | <.0001 | -13358 | -0.006 | 1.000 |
| EF0333 S1ca - BF706 S1wa | -18.211 | -7.674 | <.0001 | 413486 | 0.177 | 1.000 |
| EF0333 S1ca - EF0738 S1wa | -15.027 | -6.384 | <.0001 | 1507 | 0.001 | 1.000 |
| EF0333 S1ca - BF708 S1wa | -10.109 | -4.224 | 0.0021 | -14249 | -0.006 | 1.000 |
| EF0333 S1ca - BF709 S1wa | 24.963 | 10.687 | <.0001 | -11805 | -0.005 | 1.000 |
| BF706 S1wa - EF0738 S1wa | 3.184 | 1.353 | 0.9802 | 411979 | 0.177 | 1.000 |
| BF706 S1wa - BF708 S1wa | 8.102 | 3.384 | 0.0429 | 427735 | 0.183 | 1.000 |
| BF706 S1wa - BF709 S1wa | 43.174 | 18.486 | <.0001 | 425291 | 0.182 | 1.000 |
| EF0738 S1wa - BF708 S1wa | 4.918 | 2.071 | 0.6843 | 15756 | 0.007 | 1.000 |
| EF0738 S1wa - BF709 S1wa | 39.990 | 17.267 | <.0001 | 13312 | 0.006 | 1.000 |
| BF708 S1wa - BF709 S1wa | 35.073 | 14.881 | <.0001 | -2444 | -0.001 | 1.000 |
